# A monogenic and fast-responding Light-Inducible Cre recombinase as a novel optogenetic switch

**DOI:** 10.1101/2020.06.04.132548

**Authors:** Hélène Duplus-Bottin, Martin Spichty, Gérard Triqueneaux, Christophe Place, Philippe Emmanuel Mangeot, Théophile Ohlmann, Franck Vittoz, Gaël Yvert

**Author notes:** corresponding author. Contact Information: Gael Yvert, Laboratory of Biology and Modeling of the Cell, Ecole Normale Supérieure de Lyon, CNRS, Université de Lyon, 46 Allée d’Italie, Lyon, F-69007, France. Laboratoire d’Innovation Moléculaire et Applications, Site de Mulhouse – IRJBD, 3 bis rue Alfred Werner, 68057 Mulhouse Cedex, France (Martin Spichty).

## Abstract

Optogenetics enables genome manipulations with high spatiotemporal resolution, opening exciting possibilities for fundamental and applied biological research. Here, we report the development of LiCre, a novel light-inducible Cre recombinase. LiCre is made of a single flavin-containing protein comprising the asLOV2 photoreceptor domain of *Avena sativa* fused to a Cre variant carrying destabilizing mutations in its N-terminal and C-terminal domains. LiCre can be activated within minutes of illumination with blue light, without the need of additional chemicals. When compared to existing photoactivatable Cre recombinases based on two split units, LiCre displayed faster and stronger activation by light as well as a lower residual activity in the dark. LiCre was efficient both in yeast, where it allowed us to control the production of *β*-carotene with light, and in human cells. Given its simplicity and performances, LiCre is particularly suited for fundamental and biomedical research, as well as for controlling industrial bioprocesses.

## INTRODUCTION

The wealth of knowledge currently available on the molecular regulations of living systems - including humans - largely results from our ability to introduce genetic changes in model organisms. Such manipulations have been extremely informative because they can unambiguously demonstrate causal effects of molecules on phenotypes. The vast majority of these manipulations were made by first establishing a mutant individual - or line of individuals - and then studying it. This classic approach has two limitations. First, the mutation is present in all cells of the individual. This complicates the analysis of the contribution of specific cells or cell-types to the phenotypic alterations that are observed at the whole-organism level. Second, when a mutation is introduced long before the phenotypic analysis, it is possible that the organism has “adapted” to it, either via compensatory regulations or, in case of mutant lines maintained over multiple generations, by compensatory mutations.

For these reasons, other approaches relying on site-specific recombinases were developed to introduce specific mutations in a restricted number of cells of the organism, and at a specific time. For instance, the Cre/LoxP system^1,2^ consists of two manipulations: a stable insertion, in all cells, of foreign 34-bp DNA sequences called LoxP, and the expression of the Cre recombinase in some cells only, where it modifies the DNA by catalyzing recombination between the LoxP sites. The result is a mosaic animal - or plant, or colony of cells - where chromosomal DNA has been rearranged in some cells only. Cre is usually introduced via a transgene that is only expressed in the cells to be mutated. The location and orientation of LoxP sites can be chosen so that recombination generates either a deletion, an inversion or a translocation. Similar systems were developed based on other recombinases/recognition targets, such as Flp/FRT^3^ or Dre-rox^4^. To control the timing of recombination, several systems were made inducible. Tight control was obtained using recombinases that are inactive unless a chemical ligand is provided to the cells. For example, the widely-used Cre-ERT chimeric protein can be activated by 4-hydroxy-tamoxifen^5^. Other inducible systems rely on chemical-induced dimerization of two halves of the recombinase. For example, the FKBP– FRB split Cre system consists of two inactive proteins that can assemble in the presence of rapamycin to form a functional recombinase complex^6^. Similar systems were reported that rendered dimerization of the split Cre fragments dependent on phytohormones^7^. Although powerful, these systems present some caveats: ligands are not always neutral to cells and can therefore perturb the biological process under investigation; since they diffuse in tissues, the control of activation is sometimes not precise enough in space and/or time; and the cost or side-effects of chemical inducers can be prohibitive for industrial or biomedical applications.

More recently, several authors modified these dimerizing split recombinases to make them inducible by light instead of chemicals. This presents several advantages because i) light can be used with extreme spatiotemporal precision and high reproducibility, ii) when applied at low energy, it is neutral to many cell types, and iii) it is very cheap and therefore scalable to industrial processes. The dimerization systems that were used come from developments made in optogenetics, where various light, oxygen or voltage (LOV) protein domains have been used as photosensory modules to control transcription^8^, protein degradation^9^, dimerization^10–12^ or subcellular relocalization^13,14^. LOV domains belong to the Per-Arnt-Sim (PAS) superfamily found in many sensors. They respond to light via a flavin cofactor located at their center. In the *asLOV2* domain, blue light generates a covalent bond between a carbon atom of a flavin mononucleotide (FMN) cofactor and a cystein side chain of the PAS fold^15,16^, resulting in a conformational change including the unfolding of a large C-terminal *α*-helical region called the J*α* helix^17,18^. Diverse optogenetics tools have been developed by fusing LOV domains to functional proteins, in ways that made the J*α* folding/unfolding critical for activity^19^. Among these tools are several photodimerizers that proved useful to control the activity of recombinases. Taslimi *et al.*^20^ reported blue-light dependent heterodimerization of a split Cre recombinase using the CIB1-CRY2 dimerizers from the plant *Arabidopsis thaliana* and others successfully used the nMag/pMag dimerizers derived from Vivid (VVD), a protein of the fungus *Neurospora crassa*^21,22^. A third system was based on dimerizers derived from the chromophore-binding photoreceptor phytochrome B (PhyB) of *A. thaliana* and its interacting factor PIF3. In this case, red light was used for stimulation instead of blue light, but the system required the addition of an expensive chemical, the chromophore phycocyanobilin^23^.

An ideal inducible recombinase is one that ensures both low basal activity and high induced activity, that is simple to implement, cheap to use and fast to induce. All dimerizing split Cre systems have in common that two protein units must be assembled in order to form one functional Cre. Thus, the probability of forming a functional recombination synapse - which normally requires four Cre molecules - is proportional to the product of the two units’ cellular concentrations to the power of four. Split systems therefore strongly depend on the efficient expression of their two different coding sequences, as previously reported^24^. An inducible system based on a single protein may avoid this limitation. Its implementation by transgenesis would also be simpler, especially in vertebrates.

We report here the development of LiCre, a novel Light-Inducible Cre recombinase that is made of a single flavin-containing protein. LiCre can be activated within minutes of illumination with blue light, without the need of additional chemicals, and it shows extremely low background activity in absence of stimulation as well as high induced activity. Using the production of carotenoids by yeast as a case example, we show that LiCre and blue light can be combined to control metabolic switches that are relevant to the problem of metabolic burden in bioprocesses. We also report that LiCre can be used efficiently in human cells, making it suitable for biomedical research. Since LiCre offers cheap and precise spatiotemporal control of a genetic switch, it is amenable to numerous biotechnological applications, even at industrial scales.

## RESULTS

### The stabilizing N-ter and C-ter α-helices of the Cre recombinase are critical for its activity

A variety of optogenetic tools have been successfully developped based on LOV domain proteins, which possess α-helices that change conformation in response to light^25^. We reasoned that fusing a LOV domain to a helical domain of Cre that is critical for its function could generate a single protein with light-dependent recombinase activity. We searched for candidate α-helices by inspecting the structure of the four Cre units complexed with two LoxP DNA targets^26,27^ (Fig. 1a-b). Each subunit folds in two domains that bind to DNA as a clamp. Guo *et al.* initially reported that helices αA and αE of the amino-terminal domain, as well as helix αN of the C-terminal domain participate to inter-units contacts^27^. This role of helix αN was later confirmed by Ennifar *et al* ^26^. Contacts between αA and αE associate all four aminoterminal domains (Fig. 1a) and contacts involving αN lock the four carboxy-terminal domains in a cyclic manner (Fig. 1b). These helices were therefore good candidates for manipulating Cre activity. We focused on αA and αN because their location at protein extremities was convenient to design chimeric fusions.

**Figure 1.**
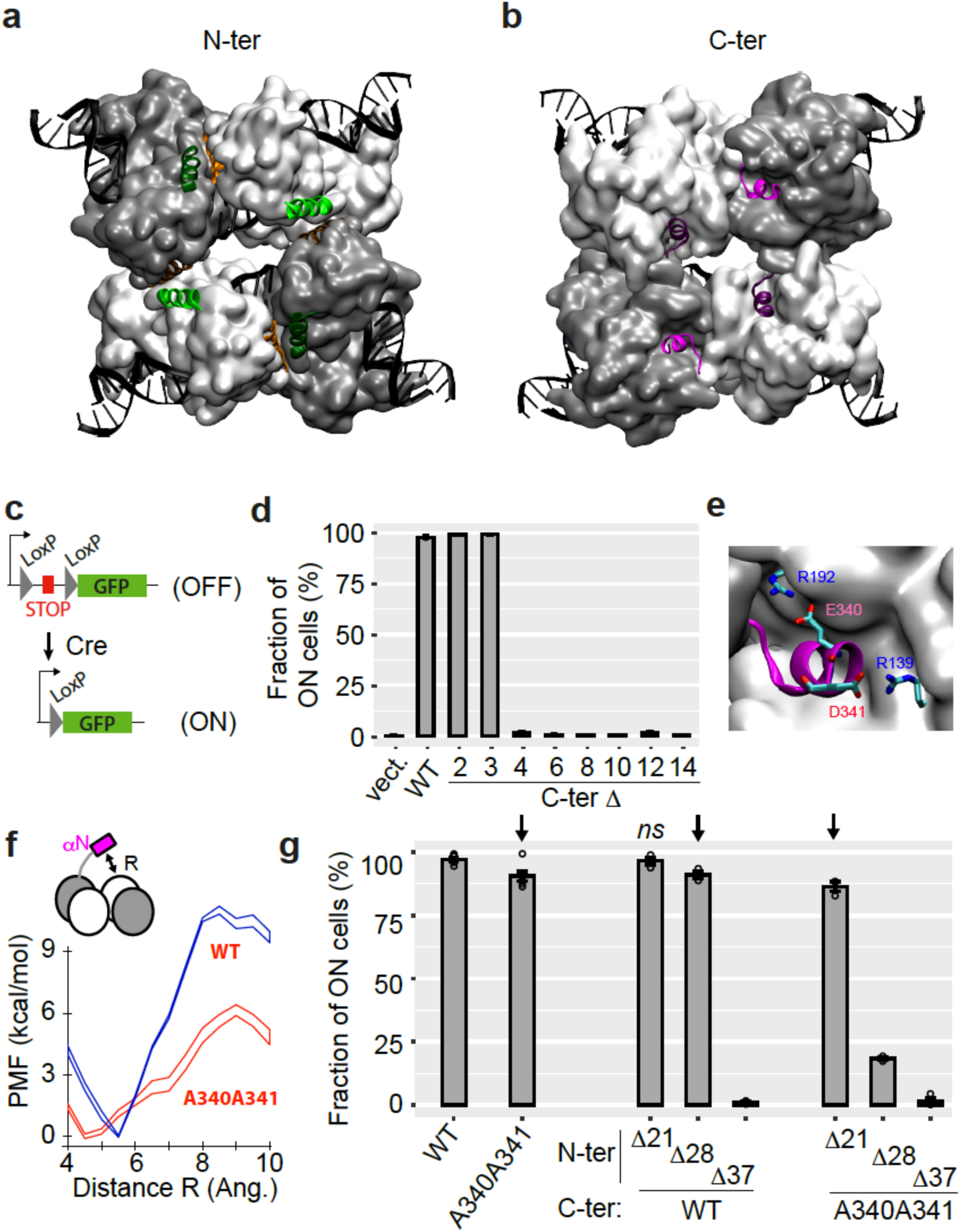
N-ter and C-ter α-helices of Cre are critical for activity. **a-b**) Structure of the Cre tetramer complexed with DNA (PDB: 1NZB). The four N-ter domains (a) interact via contacts between α-helices A (green) and E (orange) and the four C-ter domains (b) interact via α-helices N (magenta). **c**) Yeast reporter system to quantify Cre efficiency. The STOP element includes a selectable marker and a terminator sequence which prevents expression of the downstream GFP sequence. **d**) Activity of wild-type and C-ter mutants of Cre measured as the fraction of cells expressing GFP (mean +/-s.e.m, *n* = 3 independent transformants). Numbers denote the number of residues deleted from the C-ter extremity. ‘Vect’: expression plasmid with no insert. **e**) Blow-up of αN helix. **f**) Energetics of αN displacing (see methods). PMF: Potential of Mean Force (± error defined as 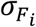 in Methods). **g**) Activity of Cre mutants lacking N-terminal residues 2 to X, combined or not with the A340 A341 C-terminal mutation (mean +/-s.e.m, *n* = 3 independent transformants). X was 21, 28 or 37 as indicated. Arrows, significantly different from WT at *p* < 0.05 (*t*-test). *ns*, non significant.

We tested the functional importance of helices αA and αN by gradually eroding them. We evaluated the corresponding mutants by expressing them in yeast cells where an active Cre can excise a repressive DNA element flanked by LoxP sites, and thereby switch ON the expression of a Green Fluorescent Protein (GFP) (Fig. 1c). After inducing the expression of Cre mutants with galactose, we counted by flow cytometry the proportion of cells that expressed GFP and we used this measure to compare recombinase activities of the different mutants (Fig. 1d). As a control, we observed that the wild-type Cre protein activated GFP expression in all cells under these conditions. Mutants lacking the last 2 or the last 3 carboxy-terminal residues displayed full activity. In contrast, mutants lacking 4 or more of the C-ter residues were totally inactive. This was consistent with a previous observation that deletion of the last 12 residues completely suppressed activity^28^. Our series of mutants showed that helix αN is needed for activity and that its residue D341 is crucial. The role of this aspartic acid is most likely to stabilize the complex: the tetramer structure indicates salt bridges between D341 and residue R139 of the adjacent unit (Fig. 1e). Interestingly, E340 might have a similar role by interacting with R192, although this residue was not essential for activity. Biomolecular simulations using a simplistic force-field model showed that the free-energy barrier for displacing the αN helix was much lower if E340 and D341 were replaced by alanines (Fig. 1f). Consistent with this prediction, we observed that a double mutant E340A D341A lost ∼10% of activity (Fig. 1g). This mild (but reproducible) reduction of activity suggested that the double mutation E340A D341A led to a fragilized version of Cre where multimerization was suboptimal.

We also tested the functional importance of α-helix A, either in a normal context where the C-terminal part of Cre was intact or where it carried the destabilizing E340A D341A mutation (Fig. 1g). Deletion of residues 2-37, which entirely ablated helix A, eliminated enzymatic activity (Fig. 1g). Very interestingly, the effect of shorter deletions depended on the C-terminal context. When the C-terminus was wild-type, removing residues 2-21 (immediately upstream of helix A) had no effect and removing residues 2-28 (partial truncation of αA) decreased the activity by ∼10%. When the C-terminus contained the E340A D341A mutation, deletions 2-21 and 2-28 were much more severe, reducing the activity by 12% and 80%, respectively. This revealed genetic interactions between the extremities of the protein, which is fully consistent with a cooperative role of helices αA and αN in stabilizing an active tetramer complex. From these observations, we considered that photo-control of Cre activity might be possible by fusing αA and αN helices to LOV domain photoreceptors.

### Fusions of LOV domains to monogenic Cre confer light-inducible activity

Our first strategy was to fuse the αN carboxy-terminal helix of Cre to the amino-terminal cap of the LOV-domain of protein Vivid (VVD), a well-characterized photosensor from *Neurospora crassa*^12,29,30^. The resulting chimeric protein, which contained the full-length Cre connected to VVD via four amino-acids, did not display light-dependent recombinase activity (Supplementary Fig. S1). Our next strategy was based on a modified version of the asLOV2 domain from *Avena sativa* which had been optimized by Guntas *et al.*^31^. These authors used it to build an optogenetic dimerizer by fusing its J*α* C-ter helix to the bacterial *SsrA* peptide. Instead, we fused J*α* to the αA amino-terminal helix of Cre. Using the same GFP reporter system as described above for detecting *in-vivo* recombination in yeast, we built a panel of constructs with various fusion positions and we directly quantified their activity with and without blue-light illumination. All fusions displayed reduced activity in both dark and light conditions as compared to wild-type Cre. Three constructs - corresponding to fusions of asLOV2 to residues 19, 27 and 32 of Cre, respectively - displayed higher activity after light stimulation. We recovered the corresponding plasmids from yeast, amplified them in bacteria to verify their sequence and re-transformed them in yeast which confirmed the differential activity between dark and light conditions for all three constructs (Fig. 2a). Fusion at position 32 (named LOV2_Cre32) displayed the highest induction by light, with activity increasing from 15% in dark condition to 50% after 30 minutes of illumination. Although this induction was significant, a 15% activity of the non-induced form remained too high for most applications. We therefore sought to reduce this residual activity, which we did in two ways.

**Figure 2.**
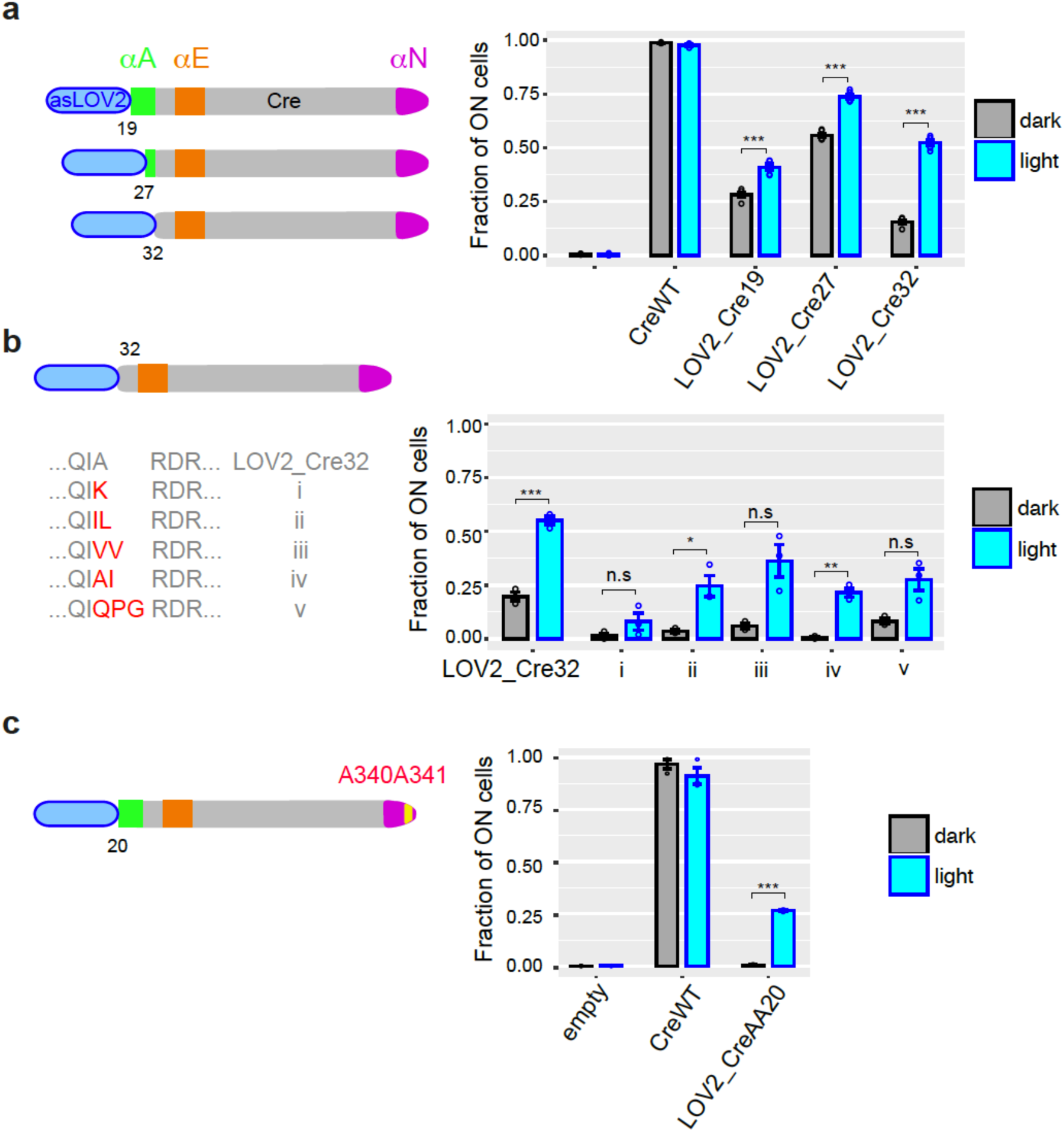
Monogenic LOV2-Cre fusions display photoactivatable recombinase activity. **a**) Fusions with wild-type Cre. **b**) Variants of LOV2_Cre32 carrying the indicated mutations at the peptide junction. **c**) Fusion with Cre carrying the A340A341 double mutation. **a -c**) Numbers indicate the positions on the Cre peptidic sequence where asLOV2 was fused. All bar plots show recombinase activity measured by flow-cytometry (mean ± sem of the proportion of switched cells, *n* = 5 independent transformants) after galactose-induced expression of the fusion protein, followed (cyan) or not (grey) by illumination at 460 nm, 36.3 mW/cm^2^, for 30 minutes. *, ** and ***: significantly different between dark and light conditions at *p* < 0.05, *p* < 0.01 and *p* < 0.001, respectively (*t*-test). *n.s.*, non significant (*p* > 0.05).

First, we randomized the residues located at the junction between asLOV2 and Cre. We used degenerate primers and *in-vivo* recombination (see methods) to mutagenize LOV2_Cre32 at these positions and we directly tested the activity of about 90 random clones. Five of them showed evidence of low residual activity in the dark and we characterized them further by sequencing and re-transformation. For all five clones, residual activity was indeed reduced as compared to LOV2_Cre32, with the strongest reduction being achieved by an isoleucine insertion at the junction position (Fig. 2b iv). However, this improvement was also accompanied by a weaker induced activity and a larger variability between independent assays.

As a complementary approach to reduce residual activity, we took advantage of the above-described genetic interaction between N-ter truncations and C-ter mutations targeting residues 340 and 341. We built another series of constructs where asLOV2 fusions to αA helix were combined with the A340A341 double mutation. This approach yielded one construct (LOV2_CreAA20), corresponding to fusion at position 20, which displayed a residual activity that was undistinguishable from the negative control, and a highly-reproducible induced activity of ∼25% (Fig. 2c). We called this construct LiCre (for ‘Light-inducible Cre’) and characterized it further.

### Efficiency and dynamics of LiCre photoactivation

We placed LiCre under the expression of the *P*_*MET17*_ promoter and we tested various illumination intensities and durations on cells that were cultured to stationary phase in absence of methionine (full expression). Activity was very low without illumination and increased with both the intensity and duration of light stimulation (Fig. 3a). The minimal intensity required for stimulation was comprised between 0.057 and 1.815 mW/cm^2^. The highest activity (∼65% of switched cells) was obtained with 90 minutes illumination at 36.3 mW/cm^2^. Extending illumination to 180 minutes did not further increase the fraction of switched cells. Remarkably, we observed that 2 minutes of illumination was enough to switch 5% of cells, and 5 minutes illumination generated 10% of switched cells (Fig. 3b).

**Figure 3.**
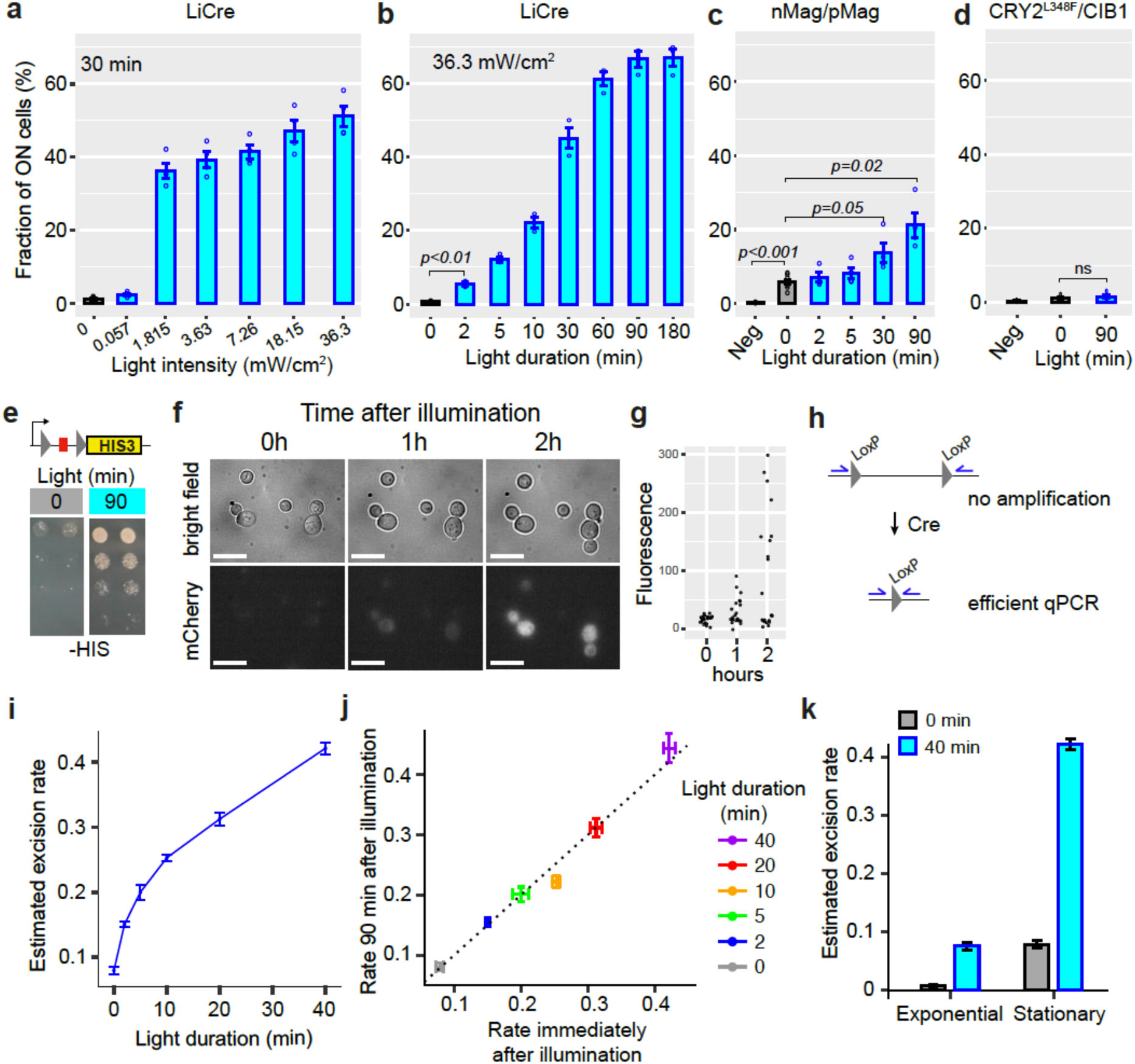
Functional properties of LiCre. **a-b**) Energy and time dependence. Yeast cells carrying the reporter system of Fig. 1c and expressing LiCre were grown to stationary phase and illuminated with blue light (460 nm, see methods) at indicated intensities, then incubated in non-dividing conditions and processed by flow cytometry (mean ± sem, strain GY1761 transformed with pGY466); *n* = 4 and 3 colonies in (a) and (b), respectively. Illumination conditions varied either in intensity (**a**) or duration (**b**). *p:* significance from *t*-test (*n*=3). The fraction of ON cells observed at 0 min was not significantly higher than the fraction of ON GY1761 cells transformed with empty vector pRS314 (*p*>0.05). **c**) Yeast strain GY1761 was transformed with plasmids pGY491 and pGY501 to express the two proteins of the nMag/pMag split Cre system of Kawano *et al.*^21^. Cells were processed as in (b) with a light intensity that matched authors recommendations (1.815 mW/cm^2^). Neg: no illumination, cells containing empty vectors only. *p:* significance from *t*-tests (*n*= 4). **d**) Yeast strain GY1761 was transformed with plasmids pGY531 and pGY532 to express the two proteins of the CRY2^L348F^/CIB1 split Cre system of Taslimi *et al.*^20^. Cells were processed as in (b) with or without illumination for 90 min at an intensity matching authors recommendations (5.45 mW/cm^2^). Neg: no illumination, cells containing empty vectors only. **e**) Yeast cells expressing LiCre from plasmid pGY466 and carrying an integrated reporter confering prototrophy to histidine were spotted on two His-plates at decreasing densities. Prior to incubation at 30**°**C, one plate (right) was illuminated for 90 minutes at 3.63 mW/cm^2^ intensity. **f**) Time-lapse imaging of yeast cells expressing LiCre and carrying a similar reporter as Fig. 1c but where GFP was replaced by mCherry (strain GY2033 with plasmid pGY466). Cells were grown to stationary phase, illuminated for 90 min at 3.63 mW/cm^2^ intensity, immobilized on bottom-glass wells in dividing condition and imaged at the indicated time. Bar: 10 µm. **g)** Quantification of intra-cellular mCherry fluorescence from (f), *n* = 22 cells. **h**) Design of qPCR assay allowing to quantify recombination efficiency. **i**) Quantification of excision by qPCR immediately after illumination at 3.63 mW/cm^2^ intensity (stationary phase, strain GY1761 transformed with pGY466). **j**) DNA excision does not occur after illumination. X-axis: same data as shown on Y-axis in (i). Y-axis: same experiment but after illumination, cells were incubated for 90 minutes in dark and non-dividing condition prior to harvest and qPCR. **k**) Quantification of DNA excision by qPCR on exponentially-growing or stationary-phase cells (strain GY1761 transformed with pGY466) illuminated at 3.63 mW/cm^2^ intensity. Grey: no illumination. Bars in (i-k): s.e.m. (*n*=10 colonies).

We compared these performances with those of two previous systems that were both based on light-dependent complementation of a split Cre enzyme. We constructed plasmids coding for proteins CreN59-nMag and pMag-CreC60 described in Kawano *et al.*^21^ and transformed them in our yeast reporter strain. Similarly, we constructed and tested plasmids coding for the proteins CRY2^L348F^-CreN and CIB1-CreC described in Taslimi *et al.*^20^. All four coding sequences were placed under the control of the yeast P_MET17_ promoter. We analyzed the resulting strains as above after adapting light to match the intensity recommended by the authors (1.815 mW/cm^2^ for nMag/pMag and 5.45 mW/cm^2^ for CRY2^L348F^/CIB1). As shown in Fig. 3c, we validated the photoactivation of nMag/pMag split Cre in yeast, where activity increased about 4-fold following 90 minutes of illumination, but we were not able to observe photoactivation of the CRY2^L348F^/CIB1 split Cre system (Fig. 3d). In addition, the photoactivation of nMag/pMag split Cre was not as fast as the one of LiCre, since 30 minutes of illumination was needed to observe a significant increase of activity. This observation is consistent with the fact that dimerization of split Cre, which is not required for LiCre, limits the rate of formation of an active recombination synapse. Another difference was that, unlike LiCre, nMag/pMag split Cre displayed a mild but significant background activity in absence of illumination (∼6% of switched cells) (Fig. 3c). Altogether, these results show that, at least in the yeast cellular context, LiCre outperforms these two other systems in terms of efficiency, rapidity and residual background activity.

To demonstrate the control of a biological activity by light, we built a reporter where Cre-mediated excision enabled the expression of the *HIS3* gene necessary for growth in absence of histidine. We cultured cells carrying this construct and expressing LiCre and we spotted them at various densities on two HIS^-^ selective plates. One plate was illuminated during 90 minutes while the other one was kept in the dark and both plates were then incubated for growth. After three days, colonies were abundant on the plate that had been illuminated and very rare on the control plate (Fig. 3e). LiCre can therefore be used to trigger cell growth with light.

We then sought to observe the switch in individual cells. To do so, we replaced GFP by mCherry in our reporter system, so that the excitation wavelength of the reporter did not overlap with stimulation of LiCre. We expressed and stimulated LiCre (90min at 3.63 mW/cm^2^) in cells carrying this reporter and subsequently imaged them over time. As expected, we observed the progressive apparition of mCherry signal in a fraction of cells (Fig. 3f-g).

Although convenient for high-throughput quantifications, reporter systems based on the *de novo* production and maturation of fluorescent proteins require a delay between the time of DNA excision and the time of acquisition. We wished to bypass this limitation and directly quantify DNA recombination. For this, we designed oligonucleotides outside of the region flanked by LoxP sites. The hybridization sites of these primers are too distant for efficient amplification of the non-edited DNA template but, after Cre-mediated excision of the internal region, these sites become proximal and PCR amplification is efficient (Fig. 3h). We mixed known amounts of edited and non-edited genomic DNA and performed real-time qPCR to build a standard curve that could be used to infer the proportion of edited DNA from qPCR signals. After this calibration, we applied this qPCR assay on genomic DNA extracted from cells collected immediately after different durations of illumination at moderate intensity (3.63 mW/cm2). Results were in full agreement with GFP-based quantifications (Fig. 3i). Excision of the target DNA occurred in a significant fraction of cells after only 2 minutes of illumination, and we estimated that excision occurred in about 30% and 40% of cells after 20 and 40 minutes of illumination, respectively. To determine if DNA excision continued to occur after switching off the light, we re-incubated half of the cells for 90 minutes in the dark prior to harvest and genomic DNA extraction. The estimated frequency of DNA excision was strikingly similar to the one measured immediately after illumination (Fig. 3j). We conclude that the reversal of activated LiCre to its inactive state is very rapid in the dark (within minutes).

The qPCR assay also allowed us to compare the efficiency of light-induced recombination between cell populations in exponential growth or in stationary phase. This revealed that LiCre photoactivation was about 4-fold more efficient in non-dividing cells (Fig. 3k). Although the reasons for this difference remain to be determined, this increase of LiCre photoactivation at stationary phase makes it particularly suitable for bioproduction applications, where metabolic switching is often desired after the growth phase (see discussion).

### Model of LiCre photo-activation

We built a structural model of LiCre to conceptualize its mode of activation (Fig. 4a). We based this model on i) the available structure of the Cre tetramer complexed with its target DNA^27^, ii) the available structure of asLOV2 in its dark state^31^ and iii) knowledge that the J*α* helix of asLOV2 domains unfolds after light activation^17,18^. From this model, we hypothesize that LiCre photoactivation may occur via two synergistic effects. First, the domain asLOV2 likely prevents Cre tetramerization in the dark state simply because of its steric occupancy. The unfolding of the J*α* helix in the light state may allow asLOV2 to liberate the multimerizing interface. Second, because the J*α* helix of asLOV2 and the αA helix of Cre are immediately adjacent, it is unlikely that both of them can fold simultaneously in their native conformation. The unfolding of J*α* in the light state may therefore stimulate proper folding of αA, and thereby allow αA to bind to the adjacent Cre unit. Structural *in vitro* studies of LiCre itself will be needed to validate these predictions.

**Figure 4.**
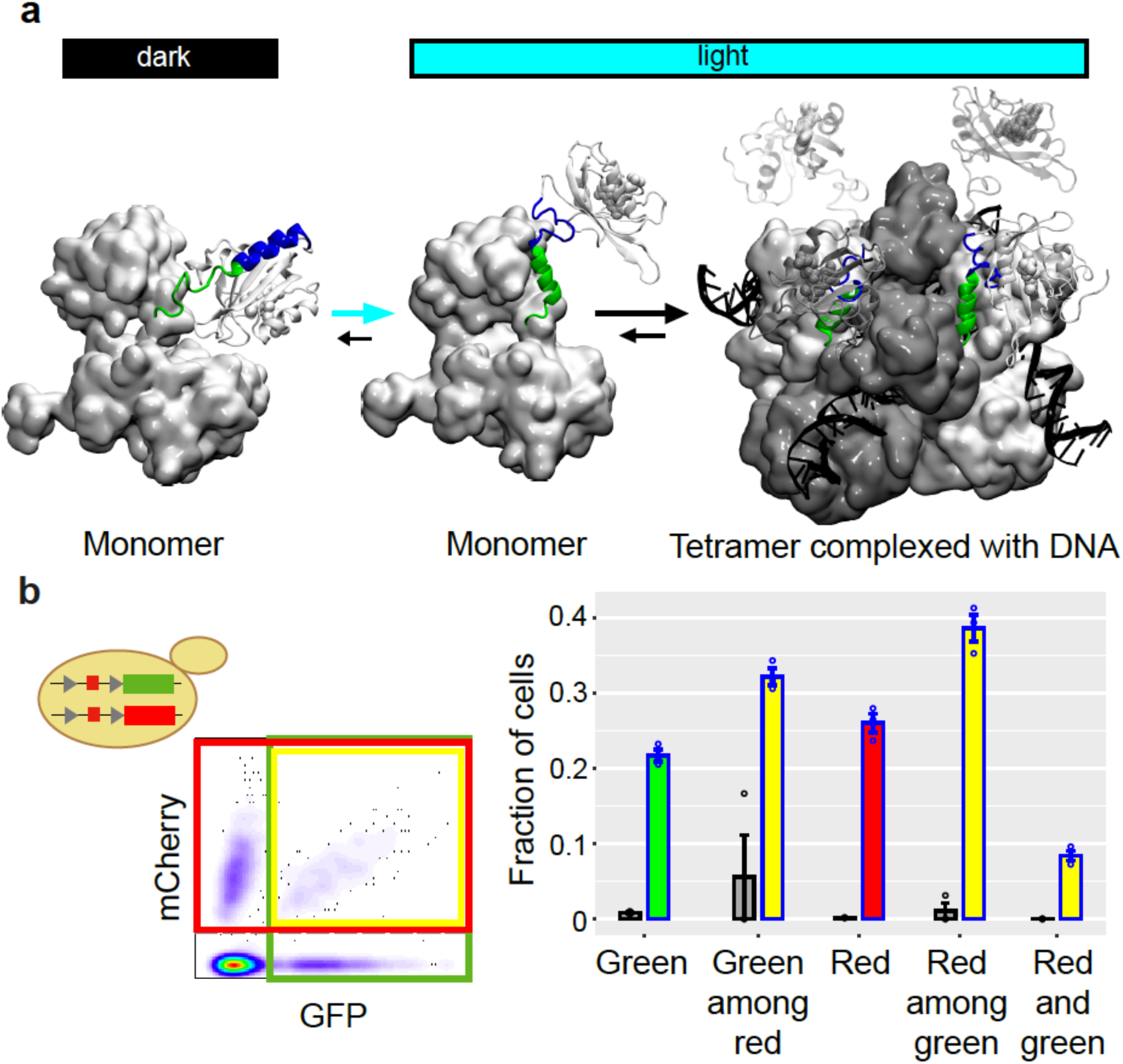
Model of LiCre activation. **a)** The model was built using PDB structures 1NZB (Cre) and 4WF0 (asLOV2). Green: residues of αA helix from Cre. Blue: residues of J*α* helix from asLOV2. **b**) Two-colors switch assay. Yeast cells carrying LiCre and both GFP and mCherry reporters (strain GY2214 with plasmid pGY466) were grown to stationary phase in SD-M-W and illuminated for 180 min at 3.63 mW/cm^2^ intensity. Cells were then incubated in the dark in non-dividing conditions and processed by flow-cytometry. Density plot (middle): fluorescent intensities for one sample. Barplot (right): mean ± s.e.m. (*n*=3 colonies) fraction of switched cells in the whole population (Green, Red, Red and green) or in the subpopulation of cells that also switched the other reporter (Green among red, Red among green). Grey bars: controls without illumination.

According to this model, there are two possible steps limiting the activation of LiCre in any one individual cell: the conformational change of LiCre monomers and the assembly of a functional recombination synapse. We sought to investigate whether one of these two steps was predominantly limiting over the other. We did this by studying cells carrying both the GFP (green) and the mCherry (red) reporters. If monomer activation is predominantly limiting, then two populations of cells are expected in an illuminated culture: cells that have activated enough LiCre molecules to form an active synapse will efficiently switch both reporters, and cells that have not activated enough LiCre monomers will leave both reporters intact and display no fluorescence. Conversely, if assembly of a functional recombination synapse is predominantly limiting, then the probability that a cell switches one reporter should be independent on what happens at the other reporter and the population will then contain a significant proportion of cells displaying fluorescence in only one color. After stimulation with 3.63 mW/cm2 blue light for 180 min, one third of the cells had switched only one of the two reporters (fluorescence in only one of the channels), ruling out the possibility that monomer activation is solely limiting (Fig. 4b). However, the probability that a reporter had switched depended on whether the other reporter had also switched. For example, the proportion of green cells in the whole population (marginal probability to switch the green reporter) was ∼20%, but the proportion of green cells in the subpopulation of red cells (conditional probability) was over 30%. Similarly, red cells were more frequent in the subpopulation of green cells than in the whole population (Fig. 4b). These observations ruled out the possibility that formation of a functional LiCre:DNA synaptic complex was solely limiting. We conclude that neither monomer activation nor synapse formation is the sole rate-limiting step *in vivo*.

### LiCre provides a light-switch for carotenoid production

LiCre offers a way to change the activities of cells without adding any chemical to their environment. This potentially makes it an interesting tool to address the limitations of metabolic burden in industrial bioproduction (see discussion). We therefore tested the possibility to use LiCre to control the production of a commercial compound with light.

Carotenoids are pigments that can be used as vitamin A precursors, anti-oxydants or coloring agents, making them valuable for the food, agriculture and cosmetics industries^32^. Commercial carotenoids are generally produced by chemical synthesis or extraction from vegetables, but alternative productions based on microbial fermentations offer remarkable advantages, including the use of low-cost substrates and therefore a high potential for financial gains. Bioproduction of carotenoids from microbes has therefore received an increasing interest. It can be based on microorganisms that naturally produce carotenoids^32^. It is also possible to introduce recombinant biosynthesis pathways in host microorganisms, which offers the advantage of a well-known physiology of the host and of optimizations by genetic engineering. For these reasons, strategies were previously developed to produce carotenoids in the yeast *S. cerevisiae*. Expressing three enzymes (*crtE, crtI* and *crtYB*) from *Xanthophyllomyces dendrorhous* enabled *S. cerevisiae* to efficiently convert farnesyl pyrophosphate (FPP) into *β*-carotene^33^. FPP is naturally produced by *S. cerevisiae* from Acetyl-CoA and serves as an intermediate metabolite, in particular for the production of ergosterol which is essential for cellular viability (Fig. 5). Thus, and as for any bioproduction consuming a cellular resource, this design is associated with a trade-off: redirecting FPP to *β*-carotene limits its availability for ergosterol biosynthesis and therefore impairs growth; and its consumption by the host cell can limit the flux towards the recombinant pathway. A promising way to deal with this trade-off would be to favor the flux towards ergosterol during biomass expansion and, after enough producer cells are obtained, to switch the demand in FPP towards *β*-carotene. We therefore explored if LiCre could offer this possibility.

**Figure 5.**
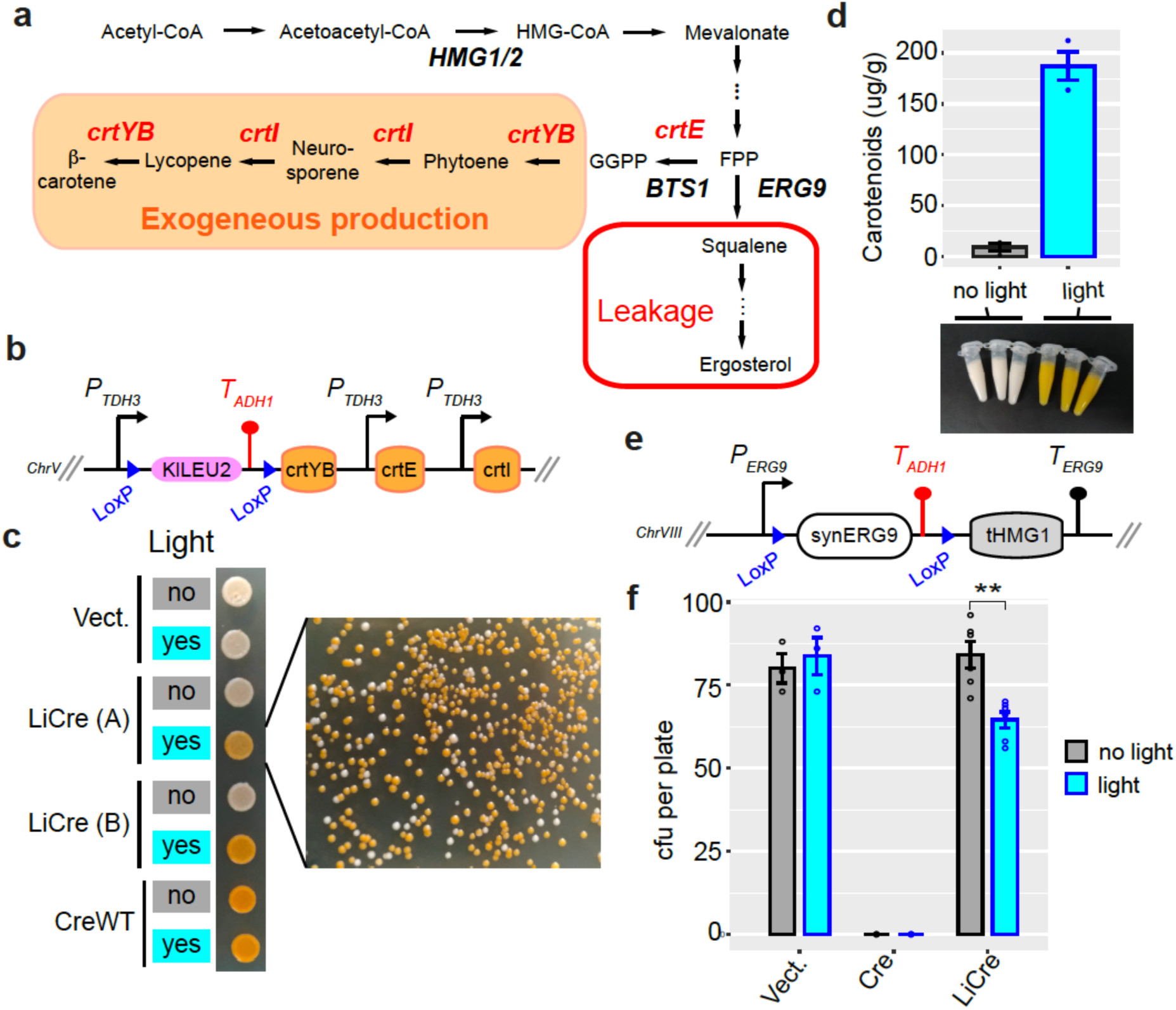
Switching ON carotenoid production with light. **a**) *β*-carotene biosynthetic pathway. Exogeneous genes from *X. dendrorhous* are printed in red. FPP, farnesyl pyrophosphate; GGPP, geranylgeranyl pyrophosphate. **b**) Scheme of the switchable locus of yeast GY2247. **c**) Photoswitchable bioproduction. Strain GY2247 was transformed with either pRS314 (Vect.), pGY466 (LiCre) or pGY502 (CreWT). Cells were cultured overnight in SD-M-W and the cultures were illuminated (460 nm, 90 min, 36.3 mW/cm^2^) or not and then spotted on agar plates. A and B correspond to two independent transformants of the LiCre plasmid. Colonies on the right originate from the illuminated LiCre (A) culture. **d**) Quantification of carotenoids production. Three colonies of strain GY2247 transformed with LiCre plasmid pGY466 were cultured overnight in SD-W-M. The following day, 10 ml of each culture were illuminated as in c), while another 10 ml was kept in the dark. These cultures were then incubated for 72 h at 30°C. Cells were pelleted (colors of the cell pelets are shown on picture) and processed for quantification (see methods). Units are micrograms of total carotenoids per gram of biomass dry weight. Bars: mean +/-s.e.m, *n* = 3. **e**) Scheme of the switchable locus of yeast GY2236. **f**) Light-induced deletion of squalene synthase gene. Strain GY2236 was transformed with either pRS314 (Vect), pGY502 (Cre) or pGY466 (LiCre). Cells were cultured overnight in 4ml of SD-M-W. A 100-µl aliquot of each culture was illuminated (as in c) while another 100-µl aliquot was kept in the dark. A dilution at ∼1 cell/µl was then plated on SD-W. Colonies were counted after 3 days. *cfu*: colony forming units (mean +/-sem, *n* ≥ 3 plates). **: *p*<0.01 (*t*-test).

First, we tested if LiCre could allow us to switch ON the exogenous production of carotenoids with light. If so, one could use it to trigger production at the desired time of a bioprocess. We constructed a *S. cerevisiae* strain expressing only two of the three enzymes required for *β*-carotene production. Expression of the third enzyme, a bifunctional phytoene synthase and lycopene cyclase, was blocked by the presence of a floxed terminator upstream of the coding sequence of the *crtYB* gene (Fig. 5b). Excision of this terminator should restore a fully-functional biosynthetic pathway. As expected, this strain formed white colonies on agar plates, but it formed orange colonies after transformation with an expression plasmid coding for Cre, indicating that *β*-carotene production was triggered (Fig. 5c). To test the possible triggering by light, we transformed this strain with a plasmid encoding LiCre and selected several transformants, which we cultured and exposed - or not - to blue light before spotting them on agar plates. The illuminated cultures became orange while the non-illuminated ones remained white. Plating a dilution of the illuminated cell suspension yielded a majority of orange colonies, indicating that LiCre triggered *crtYB* expression and *β*-carotene production in a high proportion of plated cells (Fig. 5c). We quantified bioproduction by dosing total carotenoids in cultures that had been illuminated or not. This revealed that 72 hours after the light switch the intracellular concentration of carotenoids had jumped from background levels to nearly 200µg/g (Fig. 5d). Thus, LiCre allowed us to switch ON the production of carotenoids by yeast using blue light.

We then tested if LiCre could allow us to switch OFF with light the endogenous ergosterol pathway that competes with carotenoid production for FPP consumption. The first step of this pathway is catalysed by the Erg9p squalene synthase. Given the importance of FPP availability for the production of various compounds, strategies have been reported to control the activity of this enzyme during bioprocesses, especially in order to reduce it after biomass expansion^34–36^. These strategies were not based on light but derived from transcriptional switches that naturally occur upon addition of inhibitors or when specific nutrients are exhausted from the culture medium. To test if LiCre could offer a way to switch ERG9 activity with light, we modified the *ERG9* chromosomal locus and replaced the coding sequence by a synthetic construct comprising a floxed sequence coding for Erg9p and containing a transcriptional terminator, followed by a sequence coding for the catalytic domain of the 3-hydroxy3-methylglutaryl coenzyme A reductase (tHMG1) (Fig. 5e). This design prepares *ERG9* for a Cre-mediated switch: before recombination, Erg9p is normally expressed; after recombination, ERG9 is deleted and the tHMG1 sequence is expressed to foster the mevalonate pathway. Given that ERG9 is essential for yeast viability in absence of ergosterol supplementation^37^, occurrence of the switch can be evaluated by measuring the fraction of viable yeast cells prior and after the induction of recombination. When doing so, we observed that expression of Cre completely abolished viability, regardless of illumination. In contrast, cultures expressing LiCre were highly susceptible to light: they were fully viable in absence of illumination and lost ∼23% of viable cells after light exposure (Fig. 5f). Thus, LiCre offers the possibility to abolish the activity of the yeast squalene synthase by exposing cells to light.

### LiCre switch in human cells

Beyond yeast, LiCre may also have a large spectrum of applications on multicellular organisms. Therefore, we tested its efficiency in human cells. For this, we constructed a lentiviral vector derived from the simian immunodeficiency virus (SIV) and encoding a human-optimized version of LiCre with a nuclear localization signal fused to its N-terminus. To quantify the efficiency of this vector, we also constructed a stable reporter cell line where expression of a membrane-located mCherry fluorescent protein could be switched ON by Cre/Lox recombination. We obtained this line by Flp-mediated insertion of a single copy of the reporter construct into the genome of Flp-In™ 293 cells (Fig. 6a, see methods). Our assay consisted of producing LiCre-encoding lentiviral particle, depositing them on reporter cells for 24h, illuminating the infected cultures with blue light and, 28 hours later, observing cells by fluorescence microscopy. As shown in Fig. 6b, mCherry expression was not detected in non-infected reporter cells. In cultures that were infected but not illuminated, a few positive cells were observed. In contrast, infected cultures that had been exposed to blue light contained mostly positive cells. This demonstrated the efficiency of the vector and that LiCre was poorly active unless cells were illuminated. LiCre can therefore be used to switch genetic activities in human cells with blue light.

**Figure 6.**
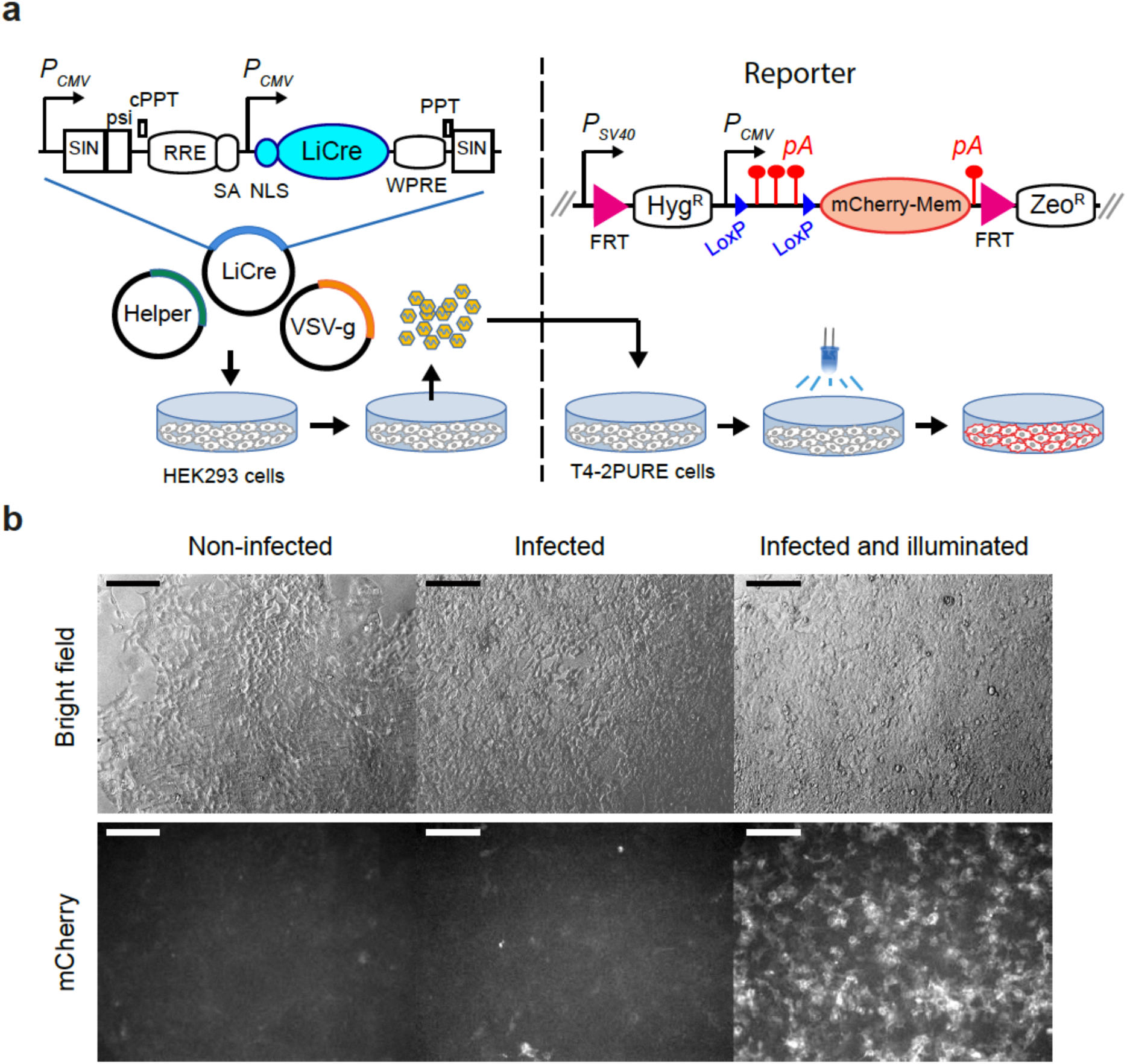
LiCre photoactivation in human cells. **a**) Left: Lentiviral SIN vector for LiCre expression (plasmid pGY577). *P*_*CMV*_, early cytomegalovirus promoter; SIN, LTR regions of Simian Immunodeficiency Virus comprising a partially-deleted 3’ U3 region followed by the R and U5 regions; *psi*, retroviral psi RNA packaging element; *cPPT* and *PPT*, central and 3’ polypurine tracks, respectively; *RRE*, Rev/Rev-responsive element; *SA*, SIV Rev/Tat splice acceptor; *NLS*, nuclear localization signal; *WPRE*, woodchuck hepatitis virus regulatory element; *Helper*, plasmid coding for *gag, pol, tat* and *rev*; *VSV-g*, plasmid encoding the envelope of the vesicular stomatitis virus. Co-transfection in HEK293T cells produces pseudotyped particles. These particles are deposited on T4-2PURE reporter cells which are then illuminated and imaged. Right, genomic reporter locus of T4-2PURE cells. *P*_*SV40*_, Promoter from SV40; *FRT*, FLP recognition targets; *Hyg*^*R*^, hygromycin resistance; *pA*, poly-adenylation signal from SV40; *Zeo*^*R*^, zeomycin resistance. Recombination between LoxP sites switches ON the expression of mCherry by removing three pA terminators. **b**) Microscopy images of T4-2PURE cells following the assay. Bars, 150 µm. All three fluorescent frames were acquired at the same intensity and exposure time. Illumination corresponded to two 20 min exposures at 3.63 mW/cm2, separated by 20 minutes without illumination.

## DISCUSSION

By performing a mutational analysis of the Cre recombinase and testing the activity of various chimeric proteins involving Cre variants and LOV-domains, we have developed a novel, single-protein, light-inducible Cre recombinase (LiCre). As compared to two previously-existing systems relying on light-dependent dimerization of split Cre fragments, LiCre displayed lower background activity in the dark as well as faster and stronger activation by light. LiCre enabled us to use blue light to switch ON the production of carotenoids by yeast and to inactivate the yeast squalene synthase. Using a lentiviral vector and human reporter cells, we also showed that LiCre could be used as an optogenetic switch in mammalian systems. We discuss below the properties of LiCre as compared to previously-reported photo-activatable recombinases and the potential of LiCre for applications in the field of industrial bioproduction.

### LiCre versus other photo-activatable recombinases

Several tools already exist for inducing site-specific recombination with light. They fall in two groups: those that require the addition of a chemical and those that are fully genetically-encoded. The first group includes the utilization of photocaged ligands instead of 4-hydroxy-tamoxifen to induce the activity of Cre-ERT. This pioneering approach was successful in cultured human cells^38^ as well as fish^39^ and mouse^40^. Later, a more complex strategy was developed that directly rendered the active site of Cre photoactivatable via the incorporation of photocaged amino-acids^41^. In this case, cells were provided with non-natural amino-acids, such as the photocaged tyrosine ONBY, and were genetically modified in order to express three foreign entities: a specifically evolved pyrrolysyl tRNA synthetase, a pyrrolysine tRNA_CUA_ and a mutant version of Cre where a critical amino-acid such as Y324 was replaced by a TAG stop codon. The tRNA synthetase/tRNA_CUA_ pair allowed the incorporation of the synthetic amino-acid in place of the nonsense mutation and the resulting enzyme was inactive unless it was irradiated with violet or ultraviolet light. This strategy successfully controlled recombination in cultured human cells^41^ and zebrafish embryos^42^. We note that it presents several caveats: its combination of chemistry and transgenes is complex to implement, the presence of the tRNA synthetase/tRNA_CUA_ pair can generate off-target artificial C-terminal tails in other proteins by bypassing natural stop codons, and violet/ultraviolet light can be harmful to cells. More recently, a radically-different chemical approach was proposed which consisted of tethering an active TAT-Cre recombinase to hollow gold nanoshells^43^. When delivered to cells in culture, these particles remained trapped in intracellular endosomes. Near-infrared photostimulation triggered activity by releasing the recombinase via nanobubble generation occurring on the particle surface. A fourth system is based on the chromophore phycocyanobilin, which binds to the PhyB receptor of *A. thaliana* and makes its interaction with PIF3 dependent on red light. Photostimulation of this interaction was used to assemble split Cre units into a functional complex in yeast^23^. A major interest of these last two systems is to offer the possibility to use red light, which is less harmful to cells than blue or violet light and better penetrates tissues. However, all these strategies require to efficiently deliver chemicals to the target cells at the appropriate time before illumination; and their underlying chemistry can be expensive, especially for applications in the context of large volumes such as industrial bioprocesses.

Other systems, such as LiCre, do not need chemical additives because they are fully genetically-encoded. To our knowledge, there are currently three such systems. One is based on the sequestration of Cre between two large photo-cleavable domains^44^. The principle of light-induced protein cleavage is very interesting but its application to Cre showed important limitations: a moderate efficiency (∼30% of ON cells after the switch), the dependence on a cellular inhibitory chaperone, and the need of violet light. The two other systems are the CRY2/CIB1 and nMag/pMag split Cre, where photo-inducible dimerizers bring together two halves of the Cre protein^20,21^. An important advantage of LiCre over these systems is that it is made of a single protein. The first benefit of this is simplicity. More efforts are needed to establish transgenic organisms expressing two open reading frames (ORFs) as compared to a single one. This is particularly true for vertebrate systems, where inserting several constructs requires additional efforts for characterizing transgene insertion sites and conducting genetic crosses. For this reason, in previous studies, the two ORFs of the split Cre system were combined in a single construct, where they were separated either by an internal ribosomal entry site or by a sequence coding a self-cleaving peptide^20,21,45^. Although helpful, these solutions have important limits: with an IRES, the two ORFs are not expressed at the same level; with a self-cleaving peptide, cleavage of the precursor protein can be incomplete, generating uncleaved products with unknown activity. This was the case for nMag/pMag split Cre in mammalian cells, where a non-cleaved form at ∼72 kDa was reported and where targeted modifications of the cleavage sequence increased both the abundance of this non-cleaved form and the non-induced activity of the system^45^. The second benefit of LiCre being a single protein is to avoid problems of suboptimal stoichiometry between the two protein units, which was reported as a possible issue for CRY2/CIB1 split Cre^24^. A third benefit is to avoid possible intra-molecular recombination between the homologous parts of the two coding sequences. Although not demonstrated, this undesired possibility was suspected for nMag/pMag split Cre because its two dimerizers derive from the same sequence^45^. The other advantages of LiCre are its performances. In the present study, we used a yeast-based assay to compare LiCre with split Cre systems. Unexpectedly, although we used the improved version of the CRY2/CIB1 split Cre containing the CRY2-L348F mutation^20^, it did not generate photo-inducible recombination in our assay. This is unlikely due to specificities of the budding yeast, such as improper protein expression or maturation, because the original authors reported activity in this organism^20^. We do not explain this result but it is consistent with the observations of Kawano *et al.*^21^ who detected extremely low photoactivation of the original version of the CRY2/CIB1 split Cre, and with the observations of Morikawa *et al.*^45^ who reported that the induced activity of the CRY2-L348F/CIB1 system was low and highly variable. In contrast, we validated the efficiency of nMag/pMag split Cre and so did other independent laboratories^7,46,47,45^. LiCre, however, displayed weaker residual activity than nMag/pMag split Cre in the dark. Reducing non-induced activity is essential for many applications where recombination is irreversible. Very recently, the nMag/pMag split Cre system was expressed in mice as a transgene -dubbed PA-Cre3.0 -which comprised the promoter sequence of the chicken beta actin gene (CAG) and synonymous modifications of the original self-cleaving coding sequence. The authors reported that this strategy abolished residual activity, and they attributed this improvement to a reduction of the expression level of the transgene^45^. It will therefore be interesting to introduce LiCre in mice with a similar expression system and compare it to PA-Cre3.0. Importantly, LiCre also displayed higher induced activity and a faster response to light as compared to nMag/pMag split Cre. This strong response probably results from its simplicity, since the activation of a single protein involves fewer steps than the activation of two units that must then dimerize to become functional. In conclusion, LiCre is simpler and more efficient than previously-existing photo-activatable recombinases.

### LiCre and industrial bioproduction

With their capability to convert low-cost substrates into valuable chemicals, cultured cells have become essential actors of industrial production. However, although metabolic pathways can be rewired in favor of the desired end-product, the yields of bioprocesses have remained limited by a challenging and universal phenomenon called metabolic burden. This effect corresponds to the natural trade-off between the fitness of host cells and their efficiency at producing exogenous compounds^48^. Loss of cellular fitness is sometimes due to viability issues -*e.g.* if the end-product is toxic to the producing cells -and sometimes simply to the fact that resources are allocated to the exogenous pathway rather than to the cellular needs. Reciprocally, satisfying the cellular demands can compromise the efficiency of exogenous pathways. In the case of carotenoids production by yeast, metabolic burden was shown to be substantial^49^ (*µ*_*max*_ reduced by ∼12%). This growth defect presumably involves competition for FPP, which is consumed to produce carotenoids but which is also crucially needed by cells to synthesize ergosterol, a major constituent of their membranes^50^.

To avoid the limitations caused by metabolic burden, a desired solution is to artificially control molecular activities so that they can first be chosen to maximize biomass expansion and then be changed in favor of bioconversion. Technically, this can be achieved by adding inducers or repressors of gene expression into the cell culture, such as lactose or hormones, but these molecules are too expensive to be used at industrial scales. Current solutions therefore rely on physiological changes in gene expression that occur in host cells during the course of fermentation, especially at the end of biomass expansion^51^. For example, expression of human recombinant proteins under the yeast P_MET17_ promoter can be repressed by extracellular methionine during the growth phase and triggered later after methionine is consumed^52,53^. Although useful, such strategies relying on endogenous molecular regulations have two important caveats. Ensuring their robustness requires strict control of physiological parameters; and each strategy is specific to the host organism and fermentation conditions and is therefore not transferable. Such limitations would be alleviated if one could cheaply control an artificial and generic metabolic switch.

Using light as the inducer is attractive in this regard. It is physiologically neutral to most non-photosynthetic organisms, it is extremely cheap and it can be controlled in real-time with extreme accuracy and reproducibility. In addition, because algae are sometimes used as producers, engineers have already designed efficient ways to bring light to bioreactors of various scales^54–56^. Placing metabolic activities of producing cells under optogenetic control is therefore a promising perspective and several developments have been made in this direction. Using the EL222 optogenetic expression system, Zhao *et al.* applied a two-regimes yeast fermentation with a continuous illumination that maintained ethanol metabolism during the growth phase, followed by light pulses stimulating isobutanol production during the bioconversion phase^57^. The potential of optogenetics was also illustrated by Milias-Argeitis *et al.* who designed a feedback control of *E. coli* growth and used it to stabilize fermentation performances against perturbations^58^.

We anticipate that LiCre can provide an alternative approach because it offers the possibility to induce irreversible genetic changes by a transient exposure to light. Applying light stimulation transiently on cells could be simpler to implement than continuously controlling light conditions in a bioreactor. By constructing appropriate Lox-based circuits, genetic changes can be designed beforehand to cause the desired switch of metabolic activities. The first switch that can be beneficial is the triggering of bioproduction itself. In principle, switching ON any artificially-designed bioproduction at the appropriate time after biomass expansion can avoid the cell-growth delays caused by metabolic burden. In the results above, we used carotenoids production as an example to illustrate how LiCre can be used to trigger bioproduction by transient illumination. To explore the potential gains on production yields, proof of concept experiments can now be made using LiCre in strains, media and fermentative conditions that are relevant to industrial processes.

The other switch that is often desired after biomass expansion is a reduction of the cellular demands for metabolites that are critical precursors of the product of interest. For example, reducing the activity of the yeast Erg9p squalene synthase is beneficial when producing terpenes -and in particular carotenoids -because more FPP becomes available for the pathway of interest. Previous efforts could reduce this activity by mutagenesis^59^, replacement of the native *ERG9* promoter^60^ or destabilization of the Erg9p protein^61^. In addition, several laboratories were able to implement a dynamic switch of ERG9 activity using conventional genetic rewiring. By placing expression of *ERG9* under the control of the *P*_*MET3*_ promoter, Asadollahi *et al.*^34^ and Amiri *et al.*^62^ could repress it by adding methionine to the culture medium, thereby improving the production of sesquiterpenes and linalool, respectively. For the production of artemisinin, Paddon *et al*.^63^ used the *P*_*CTR3*_ promoter and CuSO_4_, a cheaper inhibitor than methionine. Other studies placed the expression of *ERG9* under the control of the *P*_*HXT1*_ promoter, which is repressed when glucose becomes naturally exhausted from the medium^35,36,64^. In the present study, LiCre enabled us to inactivate *ERG9* by a transient illumination. Although full inactivation of *ERG9* causes cell death and is therefore not appropriate for industrial applications, our results show that it is possible to change ERG9 activity at a desired time and using an external stimulus that is cheaper than inhibitors. Rather than full inactivation, other LiCre-based strategies can now be designed to switch from a full activity to a reduced and viable activity. For example, one could insert a weak *erg9* allele at another genomic locus of our *lox-ERG9-lox* strain, so that gene deletion is partially complemented after recombination.

Given these considerations, optogenetic switches -and LiCre in particular -may allow industries to address the issue of metabolic burden by integrating lighting devices in bioreactors and by building switchable producer cells.

In conclusion, LiCre provides a cheap, simple, low-background, highly-efficient and fast-responding way to induce site-specific recombination with light. Given that it works in both yeast and mammalian cells, it opens many perspectives from fundamental and biomedical research to industrial applications.

## METHODS

### Strains and plasmids

Plasmids, strains and oligonucleotides used in this study are listed in Supplementary Tables S1, S2 and S3 respectively. LiCre plasmids are available from the corresponding author upon request.

### Yeast reporter systems

We ordered the synthesis of sequence LoxLEULoxHIS (Supplementary Text S1) from GeneCust who cloned the corresponding BamHI fragment in plasmid pHO-poly-HO to produce plasmid pGY262. The P_TEF_-loxP-KlLEU2-STOP-loxP-spHIS5 construct can be excised from pGY262 by NotI digestion for integration at the yeast *HO* locus. This way, we integrated it in a *leu2 his3* strain, which could then switch from LEU+ his-to leu-HIS+ after Cre-mediated recombination (Fig. 4e). To construct a GFP-based reporter, we ordered the synthesis of sequence LEULoxGreen (Supplementary Text S1) from GeneCust who cloned the corresponding NheI-SacI fragment into pGY262 to obtain pGY407. We generated strain GY984 by crossing BY4726 with FYC2-6B. We transformed GY984 with the 4-Kb NotI insert of pGY407 and obtained strain GY1752. To remove the *ade2* marker, we crossed GY1752 with FYC2-6A and obtained strain GY1761. Plasmid pGY537 targeting integration at the *LYS2* locus was obtained by cloning the BamHI-EcoRI fragment of pGY407 into the BamHI, EcoRI sites of pIS385. Plasmid pGY472 was produced by GeneCust who synthesized sequence LEULoxmCherry (Supplementary Text S1) and cloned the corresponding AgeI-EcoRI insert into the AgeI,EcoRI sites of pGY407. We generated GY983 by crossing BY4725 with FYC2-6A. We obtained GY2033 by transformation of FYC2-6B with a 4-Kb NotI fragment of pGY472. We obtained GY2207 by transformation of GY983 with the same 4-Kb NotI fragment of pGY472. To generate GY2206, we linearized pGY537 with NruI digestion, transformed in strain GY855 and selected a LEU+ Lys-colony (pop-in), which we re-streaked on 5-FoA plates for vector excision by counter-selection of URA3 (pop-out)^65^. Strain GY2214 was a diploid that we obtained by mating GY2206 with GY2207.

### Yeast expression plasmids

Mutations E340A D341A were introduced by GeneCust by site-directed mutagenesis of pSH63, yielding plasmid pGY372. We generated the N-terΔ21 mutant of Cre by PCR amplification of the P_GAL1_ promoter of pSH63 using primer 1L80 (forward) and mutagenic primer 1L71 (reverse), digestion of pSH63 by AgeI and co-transformation of this truncated plasmid and amplicon in a *trp1Δ63* yeast strain for homologous recombination and plasmid rescue. We combined the N-terΔ21 and the C-ter E340A D341A mutations similarly, but with pGY372 instead of pSH63. We generated N-terΔ28 and N-terΔ37 mutants, combined or not with C-ter E340A D341A mutations, by the same procedure where we changed 1L71 by mutagenic primers 1L72 and 1L73, respectively.

To generate a Cre-VVD fusion, we designed sequence CreCVII (Supplementary Text S1) where the Cre sequence from GENBANK AAG34515.1 was fused to the VVD-M135IM165I sequence from Zoltowski *et al.*^29^ via four additional residues (GGSG). We ordered its synthesis from GeneCust, and we co-transformed it in yeast with pSH63 (previously digested by NdeI and SalI) for homologous recombination and plasmid rescue. This generated pGY286. We then noticed an unfortunate error in AAG34515.1, which reads a threonine instead of an asparagine at position 327. We cured this mutation from pGY286 by site-directed mutagenesis using primers 1J47 and 1J48, which generated pGY339 which codes for Cre-VVD described in Supplementary Fig. S1. We constructed mutant C-terΔ14 of Cre by site-directed mutagenesis of pGY286 using primers 1J49 and 1J50 which simultaneously cured the N327T mutation and introduced an early stop codon. Mutants C-terΔ2, C-terΔ4, C-terΔ6, C-terΔ8, C-terΔ10, C-terΔ12 of Cre were constructed by GeneCust who introduced early stop codons in pGY339 by site-directed mutagenesis.

To test LOV2_Cre fusions, we first designed sequence EcoRI-LovCre_chimJa-BstBI (Supplementary Text S1) corresponding to the fusion of asLOV2 with Cre via an artificial α-helix. This helix was partly identical to the J*α* helix of asLOV2 and partly identical to the αA helix of Cre. This sequence was synthesized and cloned in the EcoRI and BstBI sites of pSH63 by GeneCust, yielding pGY408. We then generated and directly tested a variety of LOV2_Cre fusions. To do so, we digested pGY408 with BsiWI and MfeI and used this fragment as a recipient vector; we amplified the Cre sequence from pSH63 using primer 1G42 as the reverse primer, and one of primers 1M42 to 1M53 as the forward primer (each primer corresponding to a different fusion position); we co-transformed the resulting amplicon and the recipient vector in strain GY1761, isolated independent transformants and assayed them with the protocol of photoactivation and flow-cytometry described below. We generated and tested a variety of LOV2_CreAA fusions by following the same procedure where plasmid pGY372 was used as the PCR template instead of pSH63. A transformant corresponding to LOV2_Cre32 and showing light-dependent activity was chosen for plasmid rescue, yielding plasmid pGY415. A transformant corresponding to LOV2_CreAA20 was chosen for plasmid rescue, yielding plasmid pGY416. Sanger sequencing revealed that the fusion sequence present in pGY416 was QID instead of QIA at the peptide junction (position 149 on LiCre sequence of Supplementary Text S2). All further experiments on LiCre were derived from the fusion protein coded by pGY416.

To introduce random residues at the peptide junction of LOV2_Cre32 (Fig. 2b), we first generated pGY417 using the same procedure as for the generation of pGY415 but with pSH47 instead of pSH63 as the PCR template so that pGY417 has a URA3 marker instead of TRP1. We then ordered primers 1N24, 1N25 and 1N26 containing degenerate sequences, we used them with primer 1F14 to amplify the Cre sequence of pSH63, we co-transformed in strain GY1761 the resulting amplicons together with a recipient vector made by digesting plasmid pGY417 with NcoI and BsiWI, and we isolated and directly tested individual transformants with the protocol of photoactivation and flow-cytometry described below. Plasmids from transformants showing evidence of reduced background were rescued from yeast and sequenced, yielding pGY459 to pGY464.

To replace the *P*_*GAL1*_ promoter of pGY416 by the *P*_*MET17*_ promoter, we digested it with SacI and SpeI, we PCR-amplified the *P*_*MET17*_ promoter of plasmid pGY8 with primers 1N95 and 1N96, and we co-transformed the two products in yeast for homologous recombination, yielding plasmid pGY466. We changed the promoter of pGY415 using exactly the same procedure, yielding plasmid pGY465. We changed the promoter of pSH63 similarly, using primer 1O83 instead of 1N96, yielding plasmid pGY502.

To express the nMag/pMag split Cre system in yeast, we designed sequence CreN-nMag-NLS-T2A-NLS-pMag-CreCpartly (Supplementary Text S1) and ordered its synthesis from GeneCust. The corresponding BglII fragment was co-transformed in yeast for homologous recombination with pGY465 previously digested with BamHI (to remove asLOV2 and part of Cre), yielding plasmid pGY488 that contained the full system. We then derived two plasmids from pGY488, each one containing one half of the split system under the control of the Met17 promoter. We obtained the first plasmid (pGY491, carrying the TRP1 selection marker) by digestion of pGY488 with SfoI and SacII and co-transformation of the resulting recipient vector with a PCR product amplified from pGY465 using primers 1O80 and 1O82. We obtained the second plasmid (pGY501, carrying the URA3 selection marker) in two steps. We first removed the pMag-CreC part of pGY488 by digestion with NdeI and SacII followed by Klenow fill-in and religation. We then changed the selection marker by digestion with PfoI and KpnI and co-transformation in yeast with a PCR product amplified from pSH47 with primers 1O77 and 1O89.

To express the CRY2^L348F^/CIB1 split Cre system in yeast, we designed sequences CIB1CreCter and CRY2CreNter and ordered their synthesis from GeneCust, obtaining plasmids pGY526 and pGY527, respectively. To obtain pGY531, we extracted the synthetic insert of pGY527 by digestion with BglII and we co-transformed it in yeast with the NdeI-BamHI fragment of pGY466 for homologous recombination. To obtain pGY532, we extracted the synthetic insert of pGY526 by digestion with BglII and we co-transformed it in yeast with the SacI-BamHI fragment of pSH47 for homologous recombination.

To build a switchable strain for carotene production, we modified EUROSCARF strain Y41388 by integrating a LoxP-KlLEU2-T_ADH1_-LoxP cassette immediately upstream the CrtYB coding sequence of the chromosomally-integrated expression cassette described by Verwaal *et al.*^33^. This insertion was obtained by transforming Y41388 with a 6.6Kb BstBI fragment from plasmid pGY559 and selecting a Leu+ transformant, yielding strain GY2247. To obtain pGY559, we first deleted the crtE and crtI genes from YEplac195-YB_E_I^33^ by MluI digestion and religation. We then linearized the resulting plasmid with SpeI and co-transformed it for recombination in a *leu2Δ* yeast strain with a PCR amplicon obtained with primers 1P74 and 1P75 and template pGY407. After Leu+ selection, the plasmid was recovered from yeast, amplified in bacteria and verified by restriction digestion and sequencing.

We used CRISPR/Cas9 to build a switchable strain for squalene synthase. We cloned the synthetic sequence gERG9 (Supplementary Text S1) in the BamHI-NheI sites of the pML104 plasmid^66^ so that the resulting plasmid (pGY553) coded for a gRNA sequence targeting *ERG9*. This plasmid was transformed in GY2226 together with a repair-template corresponding to a 4.2-Kb EcoRI fragment of pGY547 that contained LoxP-synERG9-T_ADH1_-LoxP with homologous flanking sequences. The resulting strain was then crossed with Y41388 to obtain GY2236.

### Yeast culture media

We used synthetic (S) media made of 6.7 g/L Difco Yeast Nitrogen Base without Amino Acids and 2 g/L of a powder which was previously prepared by mixing the following amino-acids and nucleotides: 1 g of Adenine, 2 g of Uracil, 2 g of Alanine, 2 g of Arginine, 2 g of Aspartate, 2 g of Asparagine, 2 g of Cysteine, 2 g of Glutamate, 2 g of Glutamine, 2 g of Glycine, 2 g of Histidine, 2 g of Isoleucine, 4 g of Leucine, 2 g of Lysine, 2 g of Methionine, 2 g of Phenylalanine, 2 g of Proline, 2 g of Serine, 2 g of Threonine, 2 g of Tryptophane, 2 g of Tyrosine and 2 g of Valine. For growth in glucose condition, the medium (SD) also contained 20 g/L of D-glucose. For growth in galactose condition (induction of P_GAL1_ promoter), we added 2% final (20 g/L) raffinose and 2% final (20 g/L) galactose (SGalRaff medium). Media were adjusted to pH=5.8 by addition of NaOH 1N before autoclaving at 0.5 Bar. For auxotrophic selections or *P*_*MET17*_ induction, we used media where one or more of the amino-acids or nucleotides were omitted when preparing S. For example, SD-W-M was made as SD but without any tryptophane or methionine in the mix powder.

### Photoactivation and flow-cytometry quantification of recombinase activity

Unless mentioned otherwise, quantitative tests were done by flow-cytometry using yeast reporter strain GY1761. For photoactivation, we used a PAUL apparatus from GenIUL equipped with 460 nm blue LEDs. Using a NovaII photometer (Ophir^®^ Photonics), we measured that a 100% intensity on this apparatus corresponded to an energy of 36.3 mW/cm^2^. We used Zomei ND filters when we needed to obtain intensities that were not tunable on the device. The yeast reporter strain was transformed with the plasmid of interest, pre-cultured overnight in selective medium corresponding to conditions of transcriptional activation of the plasmid-borne Cre construct (SGalRaff-W for P_GAL1_ plasmids, SD-W-M for P_MET17_ plasmids, SD-W-U-M for split Cre systems) with no particular protection against ambient light. The saturated culture was transferred to two 96-well polystyrene flat-bottom Falcon^®^ sterile plates (100 µl per well) and one plate was illuminated at the indicated intensities while the other plate was kept in the dark. After the indicated duration of illumination, cells from the two plates were transferred to a fresh medium allowing expression of GFP but not cell division (SD-W-H or SD-W-U-H, strain GY1761 being auxotroph for histidine) and these cultures were incubated at 30°C for 90 minutes. Cells were then either analyzed immediately by flow cytometry, or blocked in PBS + 1mM sodium azide and analyzed the following day.

We acquired data for 10,000 events per sample using a FACSCalibur (BD Biosciences) or a MACSQuant VYB (Miltenyi Biotech) cytometer, after adjusting the concentration of cells in PBS. We analyzed raw data files in the R statistical environment (www.r-project.org) using custom-made scripts based on the flowCore package^70^ from bioconductor (www.bioconductor.org). We gated cells automatically by computing a perimeter of (FSC-H, SSC-H) values that contained 40% of events (using 2D-kernel density distributions). A threshold of fluorescent intensity (GFP or mCherry) was set to distinguish ON and OFF cells (*i.e.* expressing or not the reporter). To do this, we included in every experiment a negative control made of the reporter strain transformed with an empty vector, and we chose the 99.9^th^ percentile of the corresponding 4,000 fluorescent values (gated cells) as the threshold.

### Quantification of fluorescence levels from microscopy images

For Fig. 3g, we segmented individual cells on bright field images using the ImageJ Lasso plugin. Then, we measured on the fluorescence images the mean gray value of pixels in each segmented area, providing single-cell measures of fluorescence. For each image, the background fluorescence level was quantified from eight random regions outside of cells and with areas similar to single cells. This background level was subtracted from the fluorescence level of each cell.

### Quantification of carotenoids from yeast

We strictly followed the procedure described in Verwaal *et al.*^33^, which consists of mechanical cell lysis using glass beads, addition of pyrogallol, KOH-based saponification, and extraction of carotenoids in hexane. Quantification was estimated by optical absorption at 449 nm using a Biowave spectrophotometer.

### Human reporter cell line

We built a reporter construct for Cre-mediated recombination in human cells based on Addgene’s plasmids 55779, containing a membrane-addressed mCherry sequence^67^ (mCherry-Mem) and 51269, containing a zsGreen-based reporter of Cre recombination^68^. Re-sequencing revealed that 51269 did not contain three terminator sequences but only one between the LoxP sites. We applied a multi-steps procedure to i) restore three terminators, ii) replace zsGreen with mCherry-Mem and iii) have the final reporter in a vector suitable for targeted single-site insertion. First, we inserted a LoxP site between restriction sites NheI and HindIII of pCDNA5/FRT (Invitrogen) by annealing oligonucleotides 1O98 and 1O99, digesting and cloning this adaptor with NheI and HindIII, which yielded plasmid pGY519. Second, we replaced in two different ways the zsGreen sequence of 51269 by the mCherry-Mem sequence of 55779: either by cloning a SmaI-NotI insert from 55779 into EcoRV-NotI of 51269, yielding plasmid pGY520, or by cloning a EcoRI-NotI from 55779 into EcoRI-NotI of 51269, yielding plasmid pGY521. Third, we inserted the HindIII-NotI cassette of pGY520 into the HindIII-NotI sites of pGY519, yielding plasmid pGY523. Fourth, we inserted the HindIII-NotI cassette of pGY521 into the HindIII-NotI sites of pGY519, yielding plasmid pGY524. Fifth, a HindIII-BamHI fragment of pGY523 containing one terminator, and a BglII-EcoRI fragment of pGY524 containing another terminator were simultaneously cloned as consecutive inserts in the BglII-EcoRI sites of 51269. Finally, the resulting plasmid was digested with HindIII and BamHI to produce a fragment that was cloned into the HindIII-BglII sites of pGY524 to produce pGY525.

To establish stable cell lines, Flp-In™ T-REx™ 293 cells were purchased from Invitrogen (ThermoFisher) and transfected with both the Flp recombinase vector (pOG44, Invitrogen) and pGY525. Selection of clonal cells was first performed in medium containing 300 µg hygromycin (Sigma). After two weeks, we identified foci of cell clusters, which we individualized by transferring them to fresh wells. One of these clones was cultured for three additional weeks with high concentrations of hygromycin (up to 400 µg) to remove potentially contaminating negative cells. The resulting cell line was named T4-2PURE.

### Lentivirus construct and production

A synthetic sequence was ordered from Genecust and cloned in the HindIII-NotI sites of pCDNA3.1 (Invitrogen™ V79020). This insert contained an unrelated additional sequence that we removed by digestion with BamHI and XbaI followed by blunt-ending with Klenow fill-in. The resulting plasmid (pGY561) encoded LiCre optimized for mammalian codon usage, in-frame with a N-ter located SV40-NLS signal. This NLS-LiCre sequence was amplified from pGY561 using primers Sauci and Flard (Table S3), and the resulting amplicon was cloned in the AgeI-HindIII sites of the GAE0 Self-Inactivating Vector^69^, yielding pGY577. Lentiviral particles were produced in Gesicle Producer 293T cells (TAKARA ref 632617) transiently transfected by pGY577 (40% of total DNA), an HIV-1 helper plasmid (45% of total DNA) and a plasmid encoding the VSV-g envelope (15%) as previously described^69^. Particle-containing supernatants were clarified, filtered through a 0.45-µm membrane and concentrated by ultracentrifugation at 40,000 g before resuspension in 1xPBS (100 fold concentration).

### LiCre assay in human cells

About 3×10^5^ cells of cell line T4-2PURE were plated in two 6-well plates. After 24 h, 100 µl of viral particules were added to each well. After another 24 h, one plate was illuminated with blue-light (460 nm) using the PAUL apparatus installed in a 37°C incubator, while the other plate was kept in the dark. For illumination, we applied a sequence of 20 min ON, 20 min OFF under CO_2_ atmosphere, 20 min ON where ON corresponded to 3.63 mW/cm2 illumination. Plates were then returned to the incubator and, after 28 hours, were imaged on an Axiovert135 inverted fluorescent microscope.

### Calculation of potential mean force (PMF)

We calculated the free-energy profile (reported in Figure 1f) for the unbinding of the C-terminal α-helix in the tetrameric Cre-recombinase complex^26^ (PDB Entry 1NZB) as follows. The software we used were: the CHARMM-GUI server^71^ to generate initial input files; CHARMM version c39b1^72^ to setup the structural models and subsequent umbrella sampling by molecular dynamics; WHAM, version 2.0.9 (http://membrane.urmc.rochester.edu/content/wham/) to extract the PMF; and VMD, version 1.9.2^73^ to visualize structures. To achieve sufficient sampling by molecular dynamics, we worked with a structurally reduced model system. We focused thereby only on the unbinding of the C-terminal α-helix of subunit A (residues 334:340) from subunit F. Residues that did not have at least one atom within 25 Å from residues 333 to 343 of subunit A were removed including the DNA fragments. Residues with at least one atom within 10 Å were allowed to move freely in the following simulations; the remaining residues were fixed to their positions in the crystal structure. For the calculation of the double mutant A340A341 the corresponding residues were replaced by alanine residues. The systems were simulated with the CHARMM22 force field (GBSW & CMAP parameter file) and the implicit solvation model FACTS^74^ with recommended settings for param22 (*i.e.*, cutoff of 12 Å for nonbonded interactions). Langevin dynamics were carried out with an integration time-step of 2 fs and a friction coefficient of 4 ps^-1^ for non-hydrogen atoms. The temperature of the heat bath was set to 310 K. The hydrogen bonds were constrained to their parameter values with SHAKE^75^.

The PMF was calculated for the distance between the center of mass of the α-helix (residues 334:340 of subunit A) and the center of mass of its environment (all residues that have at least one atom within 5 Å of this helix). Umbrella sampling^76^ was performed with 13 independent molecular dynamics simulations where the system was restrained to different values of the reaction coordinate (equally spaced from 4 to 10 Å) using a harmonic biasing potential with a spring constant of 20 kcal mol^-1^ Å^-1^ (GEO/MMFP module of CHARMM). Note that this module uses a pre-factor of ½ for the harmonic potential (as in the case of the program WHAM).

For each simulation the value of the reaction coordinate was saved at every time-step for 30 ns. After an equilibration phase of 5ns, we calculated for blocks of 5 ns the PMF and the probability distribution function along the reaction coordinate using the weighted histogram analysis method^77^. A total of 13 bins were used with lower and upper boundaries at 3.75 and 10.25 Å, respectively, and a convergence tolerance of 0.01 kcal/mol. Finally, we determined for each bin its relative free energy *F*_*i*_ = −*kT* In 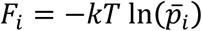 where *k* was the Boltzmann constant, *T* the temperature (310 K) and 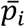 the mean value of the probability of bin *i* when averaged over the five blocks. The error in the *F*_*i*_ estimate was calculated with 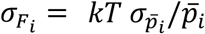 where 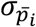 was twice the standard error of the mean of the probability. An offset was applied to the final PMF so that its lowest value was located at zero.

### qPCR quantification of recombinase activity

We grew ten colonies of strain GY1761 carrying plasmid pGY466 overnight at 30°C in SD-L-W-M liquid cultures. The following day, we used these starter cultures to inoculate 12 ml of SD-W-M medium at OD_600_ = 0.2. When monitoring growth by optical density measurements, we observed that it was fully exponential after 4 hours and until at least 8.5 hours. At 6.5 hours of growth, for each culture, we dispatched 0.1 ml in 96-well plate duplicates using one column (8 wells) per colony, we stored aliquots by centrifuging 1 ml of the cell suspension at 3300 g and freezing the cell pellet at -20°C (‘Exponential’ negative control) and we re-incubated the remaining of the culture at 30°C for later analysis at stationary phase. We exposed one plate (Fig. 4j ‘Exponential’ cyan samples) to blue light (PAUL apparatus, 460 nm, 3.63 mW/cm^2^ intensity) for 40 min while the replicate plate was kept in the dark (Fig. 4j ‘Exponential’ grey samples). We pooled cells of the same column and stored them by centrifugation and freezing as above. The following day, we collected 1 ml of each saturated, froze and stored cells as above (‘Stationary’ negative control). We dispatched the remaining of the cultures in a series of 96-well plates (0.1 ml/well, two columns per colony) and we exposed these plates to blue light (PAUL apparatus, 460 nm, 3.63 mW/cm^2^ intensity) for the indicated time (0, 2, 5, 10, 20 or 40 min). For each plate, following illumination, we collected and froze cells from 6 columns (Fig. 4h samples) and we reincubated the plate in the dark for 90 min before collecting and freezing the remaining 6 columns (Fig. 4i, x-axis samples). For genomic DNA extraction, we pooled cells from 6 wells of the same colony (1 column), we centrifuged and resuspended them in 280 µl in 50 mM EDTA, we added 20 µl of a 2 mg/ml Zymolyase stock solution (SEIKAGAKU, 20 U/mg) to the cell suspension and incubated it for 1h at 37°C for cell wall digestion. We then processed the digested cells with the Wizard Genomic DNA Purification Kit from Promega. We quantified DNA on a Nanodrop spectrophotometer and used ∼100,000 copies of genomic DNA as template for qPCR, with primers 1P57 and 1P58 to amplify the edited target and with primers 1B12, 1C22 to amplify a control HMLalpha region that we used for normalization. We ran these reactions on a Rotorgene thermocycler (Qiagen). This allowed us to quantify the rate of excision of the floxed region as N_Lox_ / N_Total_, where N_Lox_ was the number of edited molecules and N_Total_ the total number of DNA template molecules. To estimate N_Lox_, we prepared mixtures of edited and non-edited genomic DNAs, at known ratios of 0%, 0.5%, 1%, 5%, 10%, 50%, 70%, 90%, 100% and we applied (1P57,1P58) qPCR using these mixtures as templates. This provided us with a standard curve that we then used to convert Ct values of the samples of interest into N_Lox_ values. To estimate N_Total_, we qPCR-amplified the HMLalpha control region from templates made of increasing concentrations of genomic DNA. We then used the corresponding standard curve to convert the Ct value of HMLalpha amplification obtained from the samples of interest into N_Total_ values.

## ACKNOWLEDGEMENTS

We thank Fabien Duveau for critical reading of the manuscript and for signal quantifications from microscopy images, Grégory Batt for fruitful discussions, Maria Teresa Texeira for strains, Sandrine Mouradian, Véronique Barateau and SFR Biosciences Gerland-Lyon Sud (UMS344/US8) for access to flow cytometers and technical assistance, and developers of R, Bioconductor, VMD and Ubuntu for their software. This work was supported by the European Research Council under the European Union’s Seventh Framework Programme FP7/2007-2013 Grant Agreement n°281359.

## AUTHORS CONTRIBUTIONS

Constructed plasmids and strains, performed flow cytometry and yeast experiments: H.D-B.

Performed qPCR and human cell experiments: G.T.

Performed PMF computations: M.S.

Designed and produced lentiviral vector: P.M., G.T.

Conceived, designed and supervised the study, analysed flow-cytometry data: G.Y. Wrote the paper: GY.

Contributed material: C.P., F.V., T.O.

## REFERENCES

1. Duyne, G. D. V. Cre Recombinase. Microbiol. Spectr. 3, (2015).

2. Rajewsky, K. et al. Conditional gene targeting. J. Clin. Invest. 98, 600–603 (1996).

3. Lee, T. & Luo, L. Mosaic analysis with a repressible cell marker (MARCM) for Drosophila neural development. Trends Neurosci. 24, 251–254 (2001).

4. Anastassiadis, K. et al. Dre recombinase, like Cre, is a highly efficient site-specific recombinase in E. coli, mammalian cells and mice. Dis. Model. Mech. 2, 508–515 (2009).

5. Feil, R. et al. Ligand-activated site-specific recombination in mice. Proc. Natl. Acad. Sci. 93, 10887–10890 (1996).

6. Jullien, N., Sampieri, F., Enjalbert, A. & Herman, J. P. Regulation of Cre recombinase by ligand-induced complementation of inactive fragments. Nucleic Acids Res. 31, e131–e131 (2003).

7. Weinberg, B. H. et al. High-performance chemical- and light-inducible recombinases in mammalian cells and mice. Nat. Commun. 10, 4845 (2019).

8. de Mena, L., Rizk, P. & Rincon-Limas, D. E. Bringing Light to Transcription: The Optogenetics Repertoire. Front. Genet. 9, (2018).

9. Renicke, C., Schuster, D., Usherenko, S., Essen, L.-O. & Taxis, C. A LOV2 Domain-Based Optogenetic Tool to Control Protein Degradation and Cellular Function. Chem. Biol. 20, 619–626 (2013).

10. Kennedy, M. J. et al. Rapid blue-light-mediated induction of protein interactions in living cells. Nat. Methods 7, 973–975 (2010).

11. Strickland, D. et al. TULIPs: tunable, light-controlled interacting protein tags for cell biology. Nat. Methods 9, 379–384 (2012).

12. Nihongaki, Y., Suzuki, H., Kawano, F. & Sato, M. Genetically Engineered Photoinducible Homodimerization System with Improved Dimer-Forming Efficiency. ACS Chem. Biol. 9, 617–621 (2014).

13. Niopek, D. et al. Engineering light-inducible nuclear localization signals for precise spatiotemporal control of protein dynamics in living cells. Nat. Commun. 5, 1–11 (2014).

14. Witte, K., Strickland, D. & Glotzer, M. Cell cycle entry triggers a switch between two modes of Cdc42 activation during yeast polarization. eLife 6, e26722 (2017).

15. Crosson, S. & Moffat, K. Structure of a flavin-binding plant photoreceptor domain: Insights into light-mediated signal transduction. Proc. Natl. Acad. Sci. 98, 2995–3000 (2001).

16. Swartz, T. E. et al. The Photocycle of a Flavin-binding Domain of the Blue Light Photoreceptor Phototropin. J. Biol. Chem. 276, 36493–36500 (2001).

17. Swartz, T. E., Wenzel, P. J., Corchnoy, S. B., Briggs, W. R. & Bogomolni, R. A. Vibration Spectroscopy Reveals Light-Induced Chromophore and Protein Structural Changes in the LOV2 Domain of the Plant Blue-Light Receptor Phototropin 1. Biochemistry 41, 7183–7189 (2002).

18. Harper, S. M., Neil, L. C. & Gardner, K. H. Structural Basis of a Phototropin Light Switch. Science 301, 1541–1544 (2003).

19. Pudasaini, A., El-Arab, K. K. & Zoltowski, B. D. LOV-based optogenetic devices: light-driven modules to impart photoregulated control of cellular signaling. Front. Mol. Biosci. 2, (2015).

20. Taslimi, A. et al. Optimized second-generation CRY2-CIB dimerizers and photoactivatable Cre recombinase. Nat. Chem. Biol. 12, 425–430 (2016).

21. Kawano, F., Okazaki, R., Yazawa, M. & Sato, M. A photoactivatable Cre-loxP recombination system for optogenetic genome engineering. Nat. Chem. Biol. 12, 1059–1064 (2016).

22. Sheets, M. B., Wong, W. W. & Dunlop, M. J. Light-Inducible Recombinases for Bacterial Optogenetics. ACS Synth. Biol. 9, 227–235 (2020).

23. Hochrein, L., Mitchell, L. A., Schulz, K., Messerschmidt, K. & Mueller-Roeber, B. L-SCRaMbLE as a tool for light-controlled Cre-mediated recombination in yeast. Nat. Commun. 9, 1931 (2018).

24. Meador, K. et al. Achieving tight control of a photoactivatable Cre recombinase gene switch: new design strategies and functional characterization in mammalian cells and rodent. Nucleic Acids Res. 47, e97–e97 (2019).

25. Weitzman, M. & Hahn, K. M. Optogenetic approaches to cell migration and beyond. Curr. Opin. Cell Biol. 30, 112–120 (2014).

26. Ennifar, E., Meyer, J. E. W., Buchholz, F., Stewart, A. F. & Suck, D. Crystal structure of a wild-type Cre recombinase–loxP synapse reveals a novel spacer conformation suggesting an alternative mechanism for DNA cleavage activation. Nucleic Acids Res. 31, 5449–5460 (2003).

27. Guo, F., Gopaul, D. N. & van Duyne, G. D. Structure of Cre recombinase complexed with DNA in a site-specific recombination synapse. Nature 389, 40–6 (1997).

28. Rongrong, L., Lixia, W. & Zhongping, L. Effect of deletion mutation on the recombination activity of Cre recombinase. Acta Biochim Pol 52, 541–4 (2005).

29. Zoltowski, B. D., Vaccaro, B. & Crane, B. R. Mechanism-based tuning of a LOV domain photoreceptor. Nat Chem Biol 5, 827–34 (2009).

30. Vaidya, A. T., Chen, C. H., Dunlap, J. C., Loros, J. J. & Crane, B. R. Structure of a light-activated LOV protein dimer that regulates transcription. Sci Signal 4, ra50 (2011).

31. Guntas, G. et al. Engineering an improved light-induced dimer (iLID) for controlling the localization and activity of signaling proteins. Proc. Natl. Acad. Sci. 201417910 (2014) doi: 10.1073/pnas.1417910112.

32. Mata-Gómez, L. C., Montañez, J. C., Méndez-Zavala, A. & Aguilar, C. N. Biotechnological production of carotenoids by yeasts: an overview. Microb. Cell Factories 13, 12 (2014).

33. Verwaal, R. et al. High-Level Production of Beta-Carotene in Saccharomyces cerevisiae by Successive Transformation with Carotenogenic Genes from Xanthophyllomyces dendrorhous. Appl. Environ. Microbiol. 73, 4342–4350 (2007).

34. Asadollahi, M. A. et al. Production of plant sesquiterpenes in Saccharomyces cerevisiae: Effect of ERG9 repression on sesquiterpene biosynthesis. Biotechnol. Bioeng. 99, 666–677 (2008).

35. Xie, W., Ye, L., Lv, X., Xu, H. & Yu, H. Sequential control of biosynthetic pathways for balanced utilization of metabolic intermediates in Saccharomyces cerevisiae. Metab. Eng. 28, 8–18 (2015).

36. Tippmann, S., Scalcinati, G., Siewers, V. & Nielsen, J. Production of farnesene and santalene by Saccharomyces cerevisiae using fed-batch cultivations with RQ-controlled feed. Biotechnol. Bioeng. 113, 72–81 (2016).

37. Fegueur, M., Richard, L., Charles, A. D. & Karst, F. Isolation and primary structure of the ERG9 gene of Saccharomyces cerevisiae encoding squalene synthetase. Curr. Genet. 20, 365–372 (1991).

38. Link, K. H., Shi, Y. & Koh, J. T. Light activated recombination. J Am Chem Soc 127, 13088–9 (2005).

39. Sinha, D. K. et al. Photocontrol of protein activity in cultured cells and zebrafish with one- and two-photon illumination. Chembiochem 11, 653–63 (2010).

40. Lu, X. et al. Optochemogenetics (OCG) Allows More Precise Control of Genetic Engineering in Mice with CreER regulators. Bioconjug. Chem. 23, 1945–1951 (2012).

41. Luo, J. et al. Genetically encoded optical activation of DNA recombination in human cells. Chem. Commun. Camb. Engl. 52, 8529–8532 (2016).

42. Brown, W., Liu, J., Tsang, M. & Deiters, A. Cell-Lineage Tracing in Zebrafish Embryos with an Expanded Genetic Code. ChemBioChem 19, 1244–1249 (2018).

43. Morales, D. P. et al. Light-Triggered Genome Editing: Cre Recombinase Mediated Gene Editing with Near-Infrared Light. Small 14, 1800543 (2018).

44. Zhang, W. et al. Optogenetic control with a photocleavable protein, PhoCl. Nat. Methods 14, 391–394 (2017).

45. Morikawa, K. et al. Photoactivatable Cre recombinase 3.0 for in vivo mouse applications. Nat. Commun. 11, 1–11 (2020).

46. Takao, T. et al. Establishment of a tTA-dependent photoactivatable Cre recombinase knock-in mouse model for optogenetic genome engineering. Biochem. Biophys. Res. Commun. (2020) doi: 10.1016/j.bbrc.2020.03.015.

47. Allen, M. E. et al. An AND-Gated Drug and Photoactivatable Cre-loxP System for Spatiotemporal Control in Cell-Based Therapeutics. ACS Synth. Biol. 8, 2359–2371 (2019).

48. Wu, G. et al. Metabolic Burden: Cornerstones in Synthetic Biology and Metabolic Engineering Applications. Trends Biotechnol. 34, 652–664 (2016).

49. Verwaal, R. et al. Heterologous carotenoid production in Saccharomyces cerevisiae induces the pleiotropic drug resistance stress response. Yeast 27, 983–998 (2010).

50. Rest, M. E. van deret al. The plasma membrane of Saccharomyces cerevisiae: structure, function, and biogenesis. Microbiol. Rev. 59, 304–322 (1995).

51. Min, B. E., Hwang, H. G., Lim, H. G. & Jung, G. Y. Optimization of industrial microorganisms: recent advances in synthetic dynamic regulators. J. Ind. Microbiol. Biotechnol. 44, 89–98 (2017).

52. Solow, S. P., Sengbusch, J. & Laird, M. W. Heterologous Protein Production from the Inducible MET25 Promoter in Saccharomyces cerevisiae. Biotechnol. Prog. 21, 617–620 (2005).

53. Møller, T. S. B. et al. Human β-defensin-2 production from S. cerevisiae using the repressible MET17 promoter. Microb. Cell Factories 16, (2017).

54. Krujatz, F. et al. MicrOLED-photobioreactor: Design and characterization of a milliliter-scale Flat-Panel-Airlift-photobioreactor with optical process monitoring. Algal Res. 18, 225–234 (2016).

55. Hu, J.-Y. & Sato, T. A photobioreactor for microalgae cultivation with internal illumination considering flashing light effect and optimized light-source arrangement. Energy Convers. Manag. 133, 558–565 (2017).

56. Schreiber, C. et al. Growth of algal biomass in laboratory and in large-scale algal photobioreactors in the temperate climate of western Germany. Bioresour. Technol. 234, 140–149 (2017).

57. Zhao, E. M. et al. Optogenetic regulation of engineered cellular metabolism for microbial chemical production. Nature 555, 683–687 (2018).

58. Milias-Argeitis, A., Rullan, M., Aoki, S. K., Buchmann, P. & Khammash, M. Automated optogenetic feedback control for precise and robust regulation of gene expression and cell growth. Nat. Commun. 7, ncomms12546 (2016).

59. Zhuang, X. & Chappell, J. Building terpene production platforms in yeast. Biotechnol. Bioeng. 112, 1854–1864 (2015).

60. Yuan, J. & Ching, C.-B. Dynamic control of ERG9 expression for improved amorpha-4,11-diene production in Saccharomyces cerevisiae. Microb. Cell Factories 14, 38 (2015).

61. Peng, B. et al. A squalene synthase protein degradation method for improved sesquiterpene production in Saccharomyces cerevisiae. Metab. Eng. 39, 209–219 (2017).

62. Amiri, P., Shahpiri, A., Asadollahi, M. A., Momenbeik, F. & Partow, S. Metabolic engineering of <Emphasis Type=“Italic”>Saccharomyces cerevisiae</Emphasis> for linalool production. Biotechnol. Lett. 38, 503–508 (2016).

63. Paddon, C. J. et al. High-level semi-synthetic production of the potent antimalarial artemisinin. Nature 496, 528–532 (2013).

64. Scalcinati, G. et al. Dynamic control of gene expression in Saccharomyces cerevisiae engineered for the production of plant sesquitepene α-santalene in a fed-batch mode. Metab. Eng. 14, 91–103 (2012).

65. Sadowski, I., Su, T. C. & Parent, J. Disintegrator vectors for single-copy yeast chromosomal integration. Yeast 24, 447–55 (2007).

66. Laughery, M. F. et al. New Vectors for Simple and Streamlined CRISPR-Cas9 Genome Editing in Saccharomyces cerevisiae. Yeast Chichester Engl. 32, 711–720 (2015).

67. Yost, E. A., Mervine, S. M., Sabo, J. L., Hynes, T. R. & Berlot, C. H. Live Cell Analysis of G Protein β5 Complex Formation, Function, and Targeting. Mol. Pharmacol. 72, 812–825 (2007).

68. Hermann, M. et al. Binary recombinase systems for high-resolution conditional mutagenesis. Nucleic Acids Res. 42, 3894–3907 (2014).

69. Mangeot, P.-E., Cosset, F.-L., Colas, P. & Mikaelian, I. A universal transgene silencing method based on RNA interference. Nucleic Acids Res. 32, e102–e102 (2004).

70. Hahne, F. et al. flowCore: a Bioconductor package for high throughput flow cytometry. BMC Bioinformatics 10, 106 (2009).

71. Lee, J. et al. CHARMM-GUI Input Generator for NAMD, GROMACS, AMBER, OpenMM, and CHARMM/OpenMM Simulations Using the CHARMM36 Additive Force Field. J. Chem. Theory Comput. 12, 405–413 (2016).

72. Brooks, B. R. et al. CHARMM: The biomolecular simulation program. J. Comput. Chem. 30, 1545–1614 (2009).

73. Humphrey, W., Dalke, A. & Schulten, K. VMD: Visual molecular dynamics. J. Mol. Graph. 14, 33–38 (1996).

74. Haberthür, U. & Caflisch, A. FACTS: Fast analytical continuum treatment of solvation. J. Comput. Chem. 29, 701–715 (2008).

75. Ryckaert, J.-P., Ciccotti, G. & Berendsen, H. J. C. Numerical integration of the cartesian equations of motion of a system with constraints: molecular dynamics of n-alkanes. J. Comput. Phys. 23, 327–341 (1977).

76. Torrie, G. M. & Valleau, J. P. Nonphysical sampling distributions in Monte Carlo free-energy estimation: Umbrella sampling. J. Comput. Phys. 23, 187–199 (1977).

77. Kumar, S., Rosenberg, J. M., Bouzida, D., Swendsen, R. H. & Kollman, P. A. THE weighted histogram analysis method for free-energy calculations on biomolecules. I. The method. J. Comput. Chem. 13, 1011–1021 (1992).

