## Supplementary Figures, Tables and Text for "A monogenic and fast-responding Light-Inducible Cre recombinase as a novel optogenetic switch"

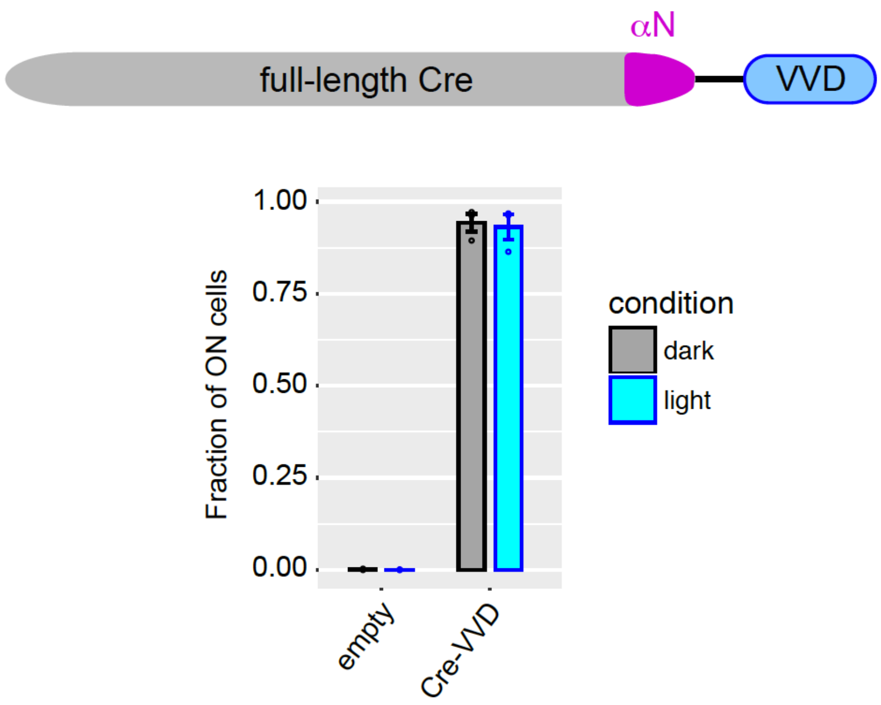

**Supplementary Figure S1: Recombinase activity of Cre-VVD chimeric protein.** Activity (fraction of switched cells) was measured after galactose-induced expression of the fusion protein, followed (cyan) or not (grey) by illumination at 460 nm, 36.3 $mW/cm^2$, for 30 minutes, followed by 90 minutes cultivation in non-dividing medium and flow-cytometry. Error bars, s.e.m (n=3 colonies of the same transformant).

Supplementary Table S1: List of plasmids used in this study.

| ID | System | Type | Selection | Description | Reference |
|---|---|---|---|---|---|
| pRS314 | yeast | CEN/ARS | TRP1 | empty vector | Sikorski et al. [1] |
| pSH47 | yeast | CEN/ARS | URA3 | pGal1:Cre | Gueldener et al. [2] |
| pSH63 | yeast | CEN/ARS | TRP1 | pGal1:Cre | Gueldener et al. [2] |
| pHO-poly-HO | yeast | Integrative | none | Vector for integration at *HO* locus | Voth et al. [3] |
| pIS385 | yeast | Integrative | URA3 | Vector for integration at *LYS2* locus | Sadowski et al.[4] |
| Addgene 51269 | human | - | - | pCAG-loxPSTOPloxP-ZsGreen reporter of Cre activity | Hermann et al.[5] |
| Addgene 55779 | human | - | NeoR | mCherry-Mem | Yost et al.[6] |
| pCDNA5/FRT | human | Integrative (FlpIn) | | Suitable for single-site insertion | Invitrogen |
| YEplac195-YB_E_I | yeast | 2-micron | URA3 | crtE-crtI-crtYB plasmid | Verwaal *et al.*[7] |
| pGY8 | yeast | Integrative | NAT | PMET17-GFP | Ansel et al. [8] |
| pGY262 | yeast | Integrative | KlLEU2 | HO-L:PTEF-loxP-KlLEU2-STOP-loxP-spHIS5:HO-R | This work |
| pGY286 | yeast | CEN/ARS | TRP1 | pGal1:Cre-VVD-N327T | This work |
| pGY339 | yeast | CEN/ARS | TRP1 | pGal1:Cre-VVD | This work |
| pGY372 | yeast | CEN/ARS | TRP1 | pGal1:CreAA | This work |
| pGY407 | yeast | Integrative | KlLEU2 | HO-L:PTEF-loxP-KlLEU2-STOP-loxP-GFP:HO-R | This work |
| pGY408 | yeast | CEN/ARS | TRP1 | pGal1:LOV2chimJaCre | This work |
| pGY415 | yeast | CEN/ARS | TRP1 | pGal1:LOV2_Cre32 | This work |
| pGY416 | yeast | CEN/ARS | TRP1 | pGal1:LiCre | This work |
| pGY417 | yeast | CEN/ARS | URA3 | pGal1:LOV2_Cre32 | This work |
| pGY459 | yeast | CEN/ARS | TRP1 | pGal1:LOV2(QIK)_Cre32 | This work |
| pGY461 | yeast | CEN/ARS | TRP1 | pGal1:LOV2(QIIL)_Cre32 | This work |
| pGY462 | yeast | CEN/ARS | TRP1 | pGal1:LOV2(QIVV)_Cre32 | This work |
| pGY463 | yeast | CEN/ARS | TRP1 | pGal1:LOV2(QIAI)_Cre32 | This work |
| pGY464 | yeast | CEN/ARS | TRP1 | pGal1:LOV2(QIQPG)_Cre32 | This work |
| pGY465 | yeast | CEN/ARS | TRP1 | pMet17:LOV2_Cre32 | This work |
| pGY466 | yeast | CEN/ARS | TRP1 | pMet17:LiCre | This work |
| pGY472 | yeast | Integrative | KlLEU2 | HO-L:PTEF-loxP-KlLEU2-STOP-loxP-mCherry:HO-R | This work |
| pGY488 | yeast | CEN/ARS | TRP1 | pMet17:Cre18-59-nMag-NLS-T2A-NLS-pMag-Cre60to343 | This work |
| pGY491 | yeast | CEN/ARS | TRP1 | pMet17:NLS-pMag-CreC60-343. | This work |
| pGY501 | yeast | CEN/ARS | URA3 | pMet17: CreN18-59-nMag-NLS | This work |

| | | | | | |
|---|---|---|---|---|---|
| pGY502 | yeast | CEN/ARS | TRP1 | pMet17:Cre | This work |
| pGY519 | human | - | - | pCDNA5 FRT loxP5 | This work |
| pGY520 | human | - | - | pCAG-Lox-STOP-MEM cherry-RV | This work |
| pGY521 | human | - | - | pCAG-Lox-STOP-MEM cherry-R1 | This work |
| pGY523 | human | - | - | pCDNA5-FRT-LoxP-1xSTOP-LoxP-MEM cherry RV | This work |
| pGY524 | human | - | - | pCDNA5-FRT-LoxP-1xSTOP-LoxP-MEM cherry R1 | This work |
| pGY525 | human | Integrative (FlpIn) | $Hyg^R$ | pcDNA5-FRT-Lox-Stop-Lox-mCherry-Mem | This work |
| pGY531 | yeast | CEN/ARS | TRP1 | pMet17:CRY2L348F-CreN19-104 | This work |
| pGY532 | yeast | CEN/ARS | URA3 | pMet17:CIB1-CreC106-343 | This work |
| pGY537 | yeast | Integrative | URA3, KlLEU2 | pISLys2:PTEF-loxP-KlLEU2-STOP-loxP-GFP | This work |
| pGY547 | yeast | Template | - | Lox-SynERG9-Lox-tHMG1 | This work |
| pGY553 | yeast | 2-micron | URA3 | pGuid-ERG9-Cas9 | This work |
| pGY559 | yeast | 2-micron | URA3 | YEplac195-PTDH3-loxP-KlLEU2-STOP-loxP-crtYB | This work |
| pGY561 | human | - | G-418 | pCDNA3.1-pCMV-NLSLiCre | This work |
| pGY577 | human | - | - | Lentiviral vector pCMV-NLS-LiCre | This work |

Supplementary Table S2: List of strains used in this study.

| ID | Background | Genotype[(i)] | Source |
|---|---|---|---|
| BY4713 | S288c (BY) | *MATb leu2Δ0* | Brachmann *et al.* [9] |
| BY4719 | S288c (BY) | *MATa trp1Δ63 ura3Δ0* | Brachmann *et al.* [9] |
| BY4725 | S288c (BY) | *MATa ade2Δ::hisG ura3Δ0* | Brachmann *et al.* [9] |
| BY4726 | S288c (BY) | *MATb ade2Δ::hisG ura3Δ0* | Brachmann *et al.* [9] |
| FYC2-6A | S288c (BY) | *MATb his3Δ200 leu2Δ1 trp1Δ63* | Gift from M.T. Teixeira |
| FYC2-6B | S288c (BY) | *MATa his3Δ200 leu2Δ1 trp1Δ63* | Gift from M.T. Teixeira |
| GY855 | S288c (BY) | *MATa leu2Δ0 trp1Δ63 ura3Δ0* | spore from BY4713 x BY4719 |
| GY983 | S288c (BY) | *MATb ade2Δ::hisG his3Δ200 leu2Δ1 trp1Δ63 ura3Δ0* | spore from BY4725 x FYC2-6A |
| GY984 | S288c (BY) | *MATa ade2Δ::hisG his3Δ200 leu2Δ1 trp1Δ63 ura3Δ0* | spore from BY4726 x FYC2-6B |
| GY1752 | S288c (BY) | *MATa ade2Δ::hisG his3Δ200 leu2Δ1 trp1Δ63 ura3Δ0 hoΔ::loxKlLEU2loxGFP* | This work |
| GY1761 | S288c (BY) | *MATb his3Δ200 leu2Δ1 trp1Δ63 ura3Δ0 hoΔ::loxKlLEU2loxGFP* | GY1752 x FYC2-6A |
| GY2033 | S288c (BY) | *MATa his3Δ200 leu2Δ1 trp1Δ63 hoΔ::loxKlLEU2loxmCherry* | This work |
| GY2206 | S288c (BY) | *MATa leu2Δ0 lys2Δ::loxKlLEU2loxGFP trp1Δ63 ura3Δ0* | This work |
| GY2207 | S288c (BY) | *MATb ade2Δ::hisG his3Δ200 leu2Δ1 trp1Δ63 ura3Δ0 hoΔ::loxKlLEU2loxmCherry* | This work |
| GY2214 | S288c (BY) | *MATa/MATb ADE2/ade2Δ::hisG his3Δ200/HIS3 leu2Δ0/leu2Δ1 LYS2/lys2Δ::loxKlLEU2loxGFP trp1Δ63/trp1Δ63 ura3Δ0/ura3Δ0 HO/hoΔ::loxKlLEU2loxmCherry* | GY2206 x GY2207 |
| GY2226 | CEN.PK | *MATb his3Δ1 leu2-3,112 trp1-289 ura3-52* | Spore from EUROSCARF strain 30000D |
| GY2236 | CEN.PK | *MATa his3Δ1 leu2-3,112 trp1-289 erg9Δ::lox-synERG9-lox-tHMG1 ura3-52::URA3-crtYB-crtI-crtE*[YIplac211-YB/I/E*]* | This work |
| GY2247 | CEN.PK | *MATa leu2-3,112 trp1-289 ura3-52::URA3-lox-KlLEU2-lox-crtYB-crtI-crtE** | This work |
| Y41388 | CEN.PK | *MATa leu2-3,112 trp1-289 ura3-52::URA3-crtYB-crtI-crtE** | EUROSCARF, from Verwaal *et al.*[7] |

(i): *MATb* corresponds to *MAT***α** *(alpha)*

Supplementary Table S3: List of DNA primers used in this study.

| ID | Sequence(i) (5'-3') |
|---|---|
| 1B12 | AAATGTCTTGTCTTCTCTGCTC |
| 1C22 | ACTGTTGCGCGAAGTAGT |
| 1F14 | gtgatgacggtgaaaacctc |
| 1G42 | AGAAAAGGAGAGGGCCAAGA |
| 1J47 | CATGAACTATATCCGTAaCCTGGATAGTGAAACAGGGGC |
| 1J48 | GCCCCTGTTTCACTATCCAGGtTACGGATATAGTTCATG |
| 1J49 | CATGAACTATATCCGTAaCCTGGATtgaGAAACAGGGGCAATGGTGCGCCTGC |
| 1J50 | GCAGGCGCACCATTGCCCCTGTTTCtcaATCCAGGtTACGGATATAGTTCATG |
| 1L71 | gaacatgtccatcaggttcttgcgaacctc-catttaacactcagataa |
| 1L72 | agaaaacgcctggcgatccctgaacatgtc-catttaacactcagataa |
| 1L73 | ggacagaagcattttccaggtatgctcaga-catttaacactcagataa |
| 1L80 | tggtatctttatagtcctgtcgg |
| 1M42 | CTT ATT AAG AAA ACC GCA TTT CAA ATC gcc gat gca acg agt gat gag gtt |
| 1M43 | CTT ATT AAG AAA ACC GCA TTT CAA ATC gcc acg agt gat gag gtt cgc aag |
| 1M44 | CTT ATT AAG AAA ACC GCA TTT CAA ATC gcc agt gat gag gtt cgc aag aac |
| 1M45 | CTT ATT AAG AAA ACC GCA TTT CAA ATC gcc gat gag gtt cgc aag aac ctg |
| 1M46 | CTT ATT AAG AAA ACC GCA TTT CAA ATC gcc gag gtt cgc aag aac ctg atg |
| 1M47 | CTT ATT AAG AAA ACC GCA TTT CAA ATC gcc gtt cgc aag aac ctg atg gac |
| 1M48 | CTT ATT AAG AAA ACC GCA TTT CAA ATC gcc cgc aag aac ctg atg gac atg |
| 1M49 | CTT ATT AAG AAA ACC GCA TTT CAA ATC gcc aag aac ctg atg gac atg ttc |
| 1M50 | CTT ATT AAG AAA ACC GCA TTT CAA ATC gcc aac ctg atg gac atg ttc agg |
| 1M51 | CTT ATT AAG AAA ACC GCA TTT CAA ATC gcc ctg atg gac atg ttc agg gat |
| 1M52 | CTT ATT AAG AAA ACC GCA TTT CAA ATC gcc gac atg ttc agg gat cgc cag |
| 1M53 | CTT ATT AAG AAA ACC GCA TTT CAA ATC gcc agg gat cgc cag gcg ttt tct g |
| 1N24 | CTT ATT AAG AAA ACC GCA TTT CAA ATC VNN AGG GAT CGC CAG GCG TTT TCT G |
| 1N25 | CTT ATT AAG AAA ACC GCA TTT CAA ATC VNN VNN AGG GAT CGC CAG GCG TTT TCT G |
| 1N26 | CTT ATT AAG AAA ACC GCA TTT CAA ATC VNN VNN VNN AGG GAT CGC CAG GCG TTT TCT G |
| 1N95 | acgccaagcgcgcaattaaccctcactaaagggaacaaaagctg-gagctc-cggatgcaagggtt |
| 1N96 | AGAGTTGTTGCTAAAGAACCGTGATGGTGGTGATGATGTGAACCTCTCAT-gtcgaggtcgacggtatcga |
| 1O77 | attaaaagatacgaggcgc |
| 1O80 | atgtgagttacctcactcat |
| 1O82 | CCAGGAGCATACAGTGTGTGACCACCGACTTTTCTCTTCTTCTTTGGTAC-CATGTCGAGGTCGACGGTATC |
| 1O83 | tcgaccggtaatgcaggcaaattttggtgtacggtcagtaaattggacat-gtcgaggtcgacggtatcga |
| 1O89 | aaggatttcaacatcgacg |
| 1P57 | CTGAGGTTCTTCTTTCATATAC |
| 1P58 | CCAGTGAATAATTCTTCACC |
| 1P74 | gttagtctttttttttagttttaaaacaccagaacttagtttcgacggattctag-TTCTCACATCACATCCGAAC |
| 1P75 | agagtatagatcagatggatctggtaatatgcgagagccgtcatggatcc- |

| | |
|---|---|
| | TTCTTCACCTTTAGAgATGG |
| Sauci | gatatcaccggtaccatgGGTCCAAAAAGAAGAGAAAGGTAGATCC |
| Flard | gatatcaagcTTAGTCGCCGGCGGCCAGCAGCCTCACCatg |

(i) N = any nucleotide, V = A, C or G.

SUPPLEMENTARY REFERENCES

1. Sikorski, R. S. & Hieter, P. A system of shuttle vectors and yeast host strains designed for efficient manipulation of DNA in Saccharomyces cerevisiae. *Genetics* **122**, 19–27 (1989).
2. Gueldener, U., Heinisch, J., Koehler, G. J., Voss, D. & Hegemann, J. H. A second set of loxP marker cassettes for Cre-mediated multiple gene knockouts in budding yeast. *Nucleic Acids Res* **30**, e23 (2002).
3. Voth, W. P., Richards, J. D., Shaw, J. M. & Stillman, D. J. Yeast vectors for integration at the HO locus. *Nucleic Acids Res* **29**, E59-9 (2001).
4. Sadowski, I., Su, T. C. & Parent, J. Disintegrator vectors for single-copy yeast chromosomal integration. *Yeast* **24**, 447–55 (2007).
5. Hermann, M. *et al.* Binary recombinase systems for high-resolution conditional mutagenesis. *Nucleic Acids Res.* **42**, 3894–3907 (2014).
6. Yost, E. A., Mervine, S. M., Sabo, J. L., Hynes, T. R. & Berlot, C. H. Live Cell Analysis of G Protein β5 Complex Formation, Function, and Targeting. *Mol. Pharmacol.* **72**, 812–825 (2007).
7. Verwaal, R. *et al.* High-Level Production of Beta-Carotene in Saccharomyces cerevisiae by Successive Transformation with Carotenogenic Genes from Xanthophyllomyces dendrorhous. *Appl. Environ. Microbiol.* **73**, 4342–4350 (2007).
8. Ansel, J. *et al.* Cell-to-cell stochastic variation in gene expression is a complex genetic trait. *PLoS Genet* **4**, e1000049 (2008).
9. Brachmann, C. B. *et al.* Designer deletion strains derived from Saccharomyces cerevisiae S288C: a useful set of strains and plasmids for PCR-mediated gene disruption and other applications. *Yeast* **14**, 115–32 (1998).

Supplementary Text S1: Synthetic nucleotidic sequences cited in Methods.

>CreCVII
GATATCCAAGCTGGTGGCTGGACCAATGTAAATATTGTCATGAACTATATCCGTACCCTGGAT
AGTGAAACAGGGGCAATGGTGCGCCTGCTGGAAGATGGCGATGGTGGTTCTGGAATGTCTCA
TACTGTTAATTCTTCTACTATGAATCCATGGGAAGTTGAAGCTTATCAACAATATCATTATG
ATCCAAGAACTGCTCCAACTGCTAATCCATTGTTCTTTCATACTTTGTATGCTCCAGGAGGTT
ATGATATTATGGGTTACCTAATTCAAATTATGAATAGACCAAATCCACAAGTGGAATTAGGT
CCCGTTGACACCTCCTGCGCCCTAATTTTATGTGACTTGAAGCAAAAAGACACTCCAATTGTG
TACGCTAGTGAAGCGTTCCTATACATGACGGGTTATTCTAATGCAGAGGTTCTGGGAAGGAA
TTGCAGGTTTTTACAGTCTCCTGACGGTATGGTGAAACCAAAATCCACAAGAAAATACGTAG
ATTCGAACACTATCAACACCATTAGGAAGGCTATCGATAGAAATGCGGAAGTACAGGTTGAA
GTCGTAAATTTCAAAAAGAACGGGCAAAGATTTGTAAACTTCTTGACCATTATTCCAGTGAG
AGATGAAACAGGTGAATATAGATACTCTATGGGTTTTCAATGTGAAACTGAAGATTATAAAG
ATGATGATGACAAATAACTCGAGTCATGTAATTAGTTATGTCACGCTTACATTCACGATATC

> CreN-nMag-NLS-T2A-NLS-pMag-CreCpartly
AGATCTCAGATACATAGATACAATTCTATTACCCCCATCCATACTCTAGAACTAGTGGATCCC
CCGGGCTGCAGGAATTCGATATCAAGCTTATCGATACCGTCGACCTCGACATGGCAACGAGTG
ATGAGGTTCGCAAGAACCTGATGGACATGTTCAGGGATCGCCAGGCGTTTTCTGAGCATACCT
GGAAAATGCTTCTGTCCGTTTGCCGGTCGTGGGCGGCATGGTGCAAGTTGAATGGGACACATA
CTTTGTACGCTCCAGGGGGCTATGACATTATGGGTTACCTGGATCAAATTGGAAATAGGCCA
AACCCTCAGGTGGAACTCGGTCCAGTAGATACTTCATGCGCTTTAATTCTTTGTGACTTAAAA
CAGAAAGATACACCTATTGTATATGCTTCCGAAGCGTTTTTGTACATGACTGGTTATTCGAAT
GCTGAAGTTCTAGGTAGGAATTGCAGATTCCTACAATCCCCTGACGGCATGGTGAAGCCTAAA
TCCACCCGAAAGTACGTCGACTCCAATACCATTAATACGATGAGAAAAGCTATTGATAGAAA
CGCTGAAGTACAAGTTGAAGTCGTCAATTTTAAGAAGAATGGCCAGAGATTCGTTAACTTCT
TGACCATGATTCCAGTAAGAGATGAAACGGGCGAATACAGGTATTCGATGGGTTTTCAATGC
GAGACTGAAGGTGGCTCCGGAGGTGTTCCGAAGAAGAAAAGAAAAGTGGGTTCCGGAGAAGG
CCGCGGTTCTCTTCTTACTTGTGGAGATGTTGAGGAAAATCCTGGTCCACTCGAGGTACCAAA
GAAGAAGAGAAAAGTCGGTGGTCACACACTGTATGCTCCTGGTGGCTATGATATCATGGGTT
ACCTAAGACAAATTAGAAATCGCCCTAATCCACAAGTCGAATTGGGTCCCGTTGATACCTCTT
GCGCTCTTATTTTGTGCGACCTAAAGCAAAAAGACACTCCTATCGTATATGCTTCGGAAGCTT
TCCTGTATATGACAGGTTACAGTAACGCCGAAGTTTTGGGGCGAAATTGCAGGTTTCTGCAA
AGCCCAGACGGGATGGTGAAGCCCAAAAGTACAAGAAAGTACGTGGACAGCAACACGATAAA
TACCATGAGAAAAGCCATTGATAGGAATGCCGAAGTACAAGTGGAAGTTGTAAACTTTAAGA
AGAATGGACAGAGATTTGTCAATTTCCTAACTATGATCCCGGTTAGGGATGAGACAGGTGAA
TACCGCTACTCGATGGGATTTCAGTGTGAAACAGAAGGCACGAACCGGAAATGGTTTCCCGCA
GAACCTGAAGATGTTCGCGATTATCTTCTATATCTTCAGGCGCGCGGTCTGGCAGTAAAAACT
ATCCAGCAACATTTGGGCCAGCTAAACATGCTTCATCGTCGGTCCGGGCTGCCACGACCAAGT
GACAGCAATGCTGTTTCACTGGTTATGCGGCGGATCCGAAAAGAAAACGTTGATGCCGGTGA
ACGTGCAAAACAGGCTCTAAGATCT

> EcoRI-LovCre_chimJa-BstBI
GAATTCGATATCAAGCTTATCGATACCGTCGAGGGGCAGAGCCGATCCTGTACACTTTACTTA
AAACCATTATCTGAGTGTTAAATGAGAGGTTCACATCATCACCACCATCACGGTTCTTTAGCA
ACAACTCTGGAAAGAATCGAGAAAAACTTCGTCATCACGGACCCGAGGCTACCCGACAACCCT
ATCATTTTCGCATCCGACTCTTTTTTACAATTAACTGAATATTCTAGAGAAGAAATTCTGGGC

CGTAATTGTAGGTTTTTACAGGGTCCAGAAACAGATAGAGCAACAGTTAGAAAGATTAGAGA
TGCTATTGATAATCAGACTGAGGTAACAGTTCAACTAATAAACTATACTAAATCCGGGAAGA
AATTTTGGAATGTCTTCCACTTACAGCCAATGAGAGATTACAAAGGTGATGTCCAGTATTTT
ATCGGAGTTCAGTTAGACGGCACAGAACGTCTGCACGGTGCCGCGGAGAGGGAAGCTGTTTG
TCTTATTAAGAAAACCGCATTTCAAATCGACGAAGCCGTACGTGACAGACAAGCATTTTCTG
AGCATACCTGGAAAATGCTTCTGTCCGTTTGCCGGTCGTGGGCGGCATGGTGCAAGTTGAATA
ACCGGAAATGGTTTCCCGCAGAACCTGAAGATGTTCGCGATTATCTTCTATATCTTCAGGCGC
GCGGTCTGGCAGTAAAAACTATCCAGCAACATTTGGGCCAGCTAAACATGCTTCATCGTCGGT
CCGGGCTGCCACGACCAAGTGACAGCAATGCTGTTTCACTGGTTATGCGGCGGATCCGAAAAG
AAAACGTTGATGCCGGTGAACGTGCAAAACAGGCTCTAGCGTTCGAA

>LoxLEULoxHIS
GGATCCGACATGGAGGCCCAGAATACCCTCCTTGACAGTCTTGACGTGCGCAGCTCAGGGGCA
TGATGTGACTGTCGCCCGTACATTTAGCCCATACATCCCCATGTATAATCATTTGCATCCATA
CATTTTGATGGCCGCACGGCGCGAAGCAAAAATTACGGCTCCTCGCTGCAGACCTGCGAGCAG
GGAAACGCTCCCCTCACAGACGCGTTGAATTGTCCCCACGCCGCGCCCCTGTAGAGAAATATA
AAAGGTTAGGATTTGCCACTGAGGTTCTTCTTTCATATACTTCCTTTTAAAATCTTGCTAGGA
TACAGTTCTCACATCACATCCGAACATAAACAACCGTTAACATAACTTCGTATAGCATACATT
ATACGAAGTTATCCATGTCTAAGAATATCGTTGTCCTACCGGGTGATCACGTCGGTAAAGAA
GTTACTGACGAAGCTATTAAGGTCTTGAATGCCATTGCTGAAGTCCGTCCAGAAATTAAGTT
CAATTTCCAACATCACTTGATCGGGGGTGCTGCCATCGATGCCACTGGCACTCCTTTACCAGA
TGAAGCTCTAGAAGCCTCTAAGAAAGCCGATGCTGTCTTACTAGGTGCTGTTGGTGGTCCAAA
ATGGGGTACGGGCGCAGTTAGACCAGAACAAGGTCTATTGAAGATCAGAAAGGAATTGGGTC
TATACGCCAACTTGAGACCATGTAACTTTGCTTCTGATTCTTTACTAGATCTTTCTCCTTTGA
AGCCTGAATATGCAAAGGGTACCGATTTCGTCGTCGTTAGAGAATTGGTTGGTGGTATCTAC
TTTGGTGAAAGAAAAGAAGATGAAGGTGACGGAGTTGCTTGGGACTCTGAGAAATACAGTGT
TCCTGAAGTTCAAAGAATTACAAGAATGGCTGCTTTCTTGGCATTGCAACAAAACCCACCATT
ACCAATCTGGTCTCTTGACAAGGCTAACGTGCTTGCCTCTTCCAGATTGTGGAGAAAGACTGT
TGAAGAAACCATCAAGACTGAGTTCCCACAATTAACTGTTCAGCACCAATTGATCGACTCTGC
TGCTATGATTTTGGTTAAATCACCAACTAAGCTAAACGGTGTTGTTATTACCAACAACATGT
TTGGTGATATTATCTCCGATGAAGCCTCTGTTATTCCAGGTTCTTTGGGTTTATTACCTTCTG
CATCTCTAGCTTCCCTACCTGACACTAACAAGGCATTCGGTTTGTACGAACCATGTCATGGTT
CTGCCCCAGATTTACCAGCAAACAAGGTTAATCCAATTGCTACCATCTTATCTGCAGCTATGA
TGTTGAAGTTATCCTTGGATTTGGTTGAAGAAGGTAGGGCTCTTGAAGAAGCTGTTAGAAAT
GTCTTGGATGCAGGTGTCAGAACCGGTGACCTTGGTGGTTCTAACTCTACCACTGAGGTTGGC
GATGCTATCGCCAAGGCTGTCAAGGAAATCTTGGCTTAACGCGCCACTTCTAAATAAGCGAAT
TTCTTATGATTTATGATTTTTATTATTAAATAAGTTATAAAAAAAATAAGTGTATACAAATT
TTAAAGTGACTCTTAGGTTTTAAAACGAAAATTCTTGTTCTTGAGTAACTCTTTCCTGTAGG
TCAGGTTGCTTTCTCAGGTATAGCATGAGGTCGCTCTTATTGACCACACCTCTACCGGCAGAT
CGCTAGCATAACTTCGTATAGCATACATTATACGAAGTTATCCATGGGTAGGAGGGCTTTTG
TAGAAAGAAATACGAACGAAACGAAAATCAGCGTTGCCATCGCTTTGGACAAAGCTCCCTTA
CCTGAAGAGTCGAATTTTATTGATGAACTTATAACTTCCAAGCATGCAAACCAAAAGGGAGA
ACAAGTAATCCAAGTAGACACGGGAATTGGATTCTTGGATCACATGTATCATGCACTGGCTA
AACATGCAGGCTGGAGCTTACGACTTTACTCAAGAGGTGATTTAATCATCGATGATCATCAC
ACTGCAGAAGATACTGCTATTGCACTTGGTATTGCATTCAAGCAGGCTATGGGTAACTTTGCC
GGCGTTAAAAGATTTGGACATGCTTATTGTCCACTTGACGAAGCTCTTTCTAGAAGCGTAGTT
GACTTGTCGGGACGGCCCTATGCTGTTATCGATTTGGGATTAAAGCGTGAAAAGGTTGGGGA
ATTGTCCTGTGAAATGATCCCTCACTTACTATATTCCTTTTCGGTAGCAGCTGGAATTACTTT
GCATGTTACCTGCTTATATGGTAGTAATGACCATCATCGTGCTGAAAGCGCTTTTAAATCTCT

GGCTGTTGCCATGCGCGCGGCTACTAGTCTTACTGGAAGTTCTGAAGTCCCAAGCACGAAGGG
AGTGTTGTAATAACTGGTCGAGTCATGTAATTAGTTATGTCACGCTTACATTCACGCCCTCCC
CCCACATCCGCTCTAACCGAAAAGGAAGGAGTTAGACAACCTGAAGTCTAGGTCCCTATTTAT
TTTTTTATAGTTATGTTAGTATTAAGAACGTTATTTATATTTCAAATTTTTCTTTTTTTTCT
GTACAGACGCGTGTACGCATGTAACATTATACTGAAAACCTTGCTTGAGAAGGTTTTGGGAC
GCTCGAAGGCTTTAATTTGCGGCCGGTACGGATCC

>LEULoxGreen
GCTAGCATAACTTCGTATAGCATACATTATACGAAGTTATCCATGTCTAAAGGTGAAGAATT
ATTCACTGGTGTTGTCCCAATTTTGGTTGAATTAGATGGTGATGTTAATGGTCACAAATTTT
CTGTCTCCGGTGAAGGTGAAGGTGATGCTACTTACGGTAAATTGACCTTAAAATTTATTTGT
ACTACTGGTAAATTGCCAGTTCCATGGCCAACCTTAGTCACTACTTTCGGTTATGGTGTTCAA
TGTTTTGCTAGATACCCAGATCATATGAAACAACATGACTTTTTCAAGTCTGCCATGCCAGAA
GGTTATGTTCAAGAAAGAACTATTTTTTTCAAAGATGACGGTAACTACAAGACCAGAGCTGA
AGTCAAGTTTGAAGGTGATACCTTAGTTAATAGAATCGAATTAAAAGGTATTGATTTTAAAG
AAGATGGTAACATTTTAGGTCACAAATTGGAATACAACTATAACTCTCACAATGTTTACATC
ATGGCTGACAAACAAAAGAATGGTATCAAAGTTAACTTCAAAATTAGACACAACATTGAAGA
TGGTTCTGTTCAATTAGCTGACCATTATCAACAAAATACTCCAATTGGTGATGGTCCAGTCTT
GTTACCAGACAACCATTACTTATCCACTCAATCTGCCTTATCCAAAGATCCAAACGAAAAGAG
AGACCACATGGTCTTGTTAGAATTTGTTACTGCTGCTGGTATTACCCATGGTATGGATGAATT
GTACAAATAACCTAGGCTGGTCGAGTCATGTAATTAGTTATGTCACGCTTACATTCACGCCCT
CCCCCCACATCCGCTCTAACCGAAAAGGAAGGAGTTAGACAACCTGAAGTCTAGGTCCCTATT
TATTTTTTTATAGTTATGTTAGTATTAAGAACGTTATTTATATTTCAAATTTTTCTTTTTTT
TCTGTACAGACGCGTGTACGCATGTAACATTATACTGAAAACCTTGCTTGAGAAGGTTTTGG
GACGCTCGAAGGCTTTAATTTGCGGCCGGTACGAGCTC

>LEULoxmCherry
ACCGGTGACCTTGGTGGTTCTAACTCTACCACTGAGGTTGGCGATGCTATCGCCAAGGCTGTC
AAGGAAATCTTGGCTTAACGCGCCACTTCTAAATAAGCGAATTTCTTATGATTTATGATTTT
TATTATTAAATAAGTTATAAAAAAAATAAGTGTATACAAATTTTAAAGTGACTCTTAGGTTT
TAAAACGAAAATTCTTGTTCTTGAGTAACTCTTTCCTGTAGGTCAGGTTGCTTTCTCAGGTAT
AGCATGAGGTCGCTCTTATTGACCACACCTCTACCGGCAGATCGCTAGCATAACTTCGTATAG
CATACATTATACGAAGTTATCCATGGTGTCAAAGGGGGAGGAAGATAATATGGCGATAATTA
AAGAGTTCATGAGGTTTAAAGTCCACATGGAGGGTTCAGTCAACGGTCATGAGTTCGAGATC
GAAGGTGAGGGTGAAGGCAGACCGTATGAAGGTACTCAAACTGCCAAATTGAAGGTGACCAA
AGGCGGCCCACTTCCGTTTGCGTGGGACATTCTTTCACCTCAATTCATGTACGGTTCGAAAGC
TTATGTTAAACATCCAGCAGATATTCCCGATTATCTAAAGTTGTCTTTCCCTGAAGGTTTTAA
ATGGGAAAGGGTCATGAATTTTGAAGACGGTGGGGTTGTAACGGTAACACAGGATTCCTCAT
TACAAGATGGCGAATTTATCTATAAAGTCAAGTTGCGTGGCACTAATTTTCCTTCTGATGGTC
CTGTCATGCAGAAGAAAACAATGGGCTGGGAAGCTTCAAGCGAAAGGATGTACCCAGAAGAT
GGTGCTTTAAAGGGTGAGATCAAACAAAGATTAAAGTTAAAGGACGGCGGGCATTACGATGC
TGAAGTTAAAACGACTTATAAAGCTAAGAAACCTGTTCAGCTGCCAGGTGCATACAACGTGA
ATATAAAGCTTGACATAACATCACATAACGAGGACTATACAATTGTTGAACAGTATGAAAGA
GCCGAAGGTCGTCACAGTACTGGAGGGATGGATGAACTATACAAATAACCTAGGCTGGTCGA
GTCATGTAATTAGTTATGTCACGCTTACATTCACGCCCTCCCCCCACATCCGCTCTAACCGAA
AAGGAAGGAGTTAGACAACCTGAAGTCTAGGTCCCTATTTATTTTTTTATAGTTATGTTAGT
ATTAAGAACGTTATTTATATTTCAAATTTTTCTTTTTTTTCTGTACAGACGCGTGTACGCATG
TAACATTATACTGAAAACCTTGCTTGAGAAGGTTTTGGGACGCTCGAAGGCTTTAATTTGCG
GCCGGTACGAGCTCGAATTC

>CIB1CreCter
TCGCGCGTTTCGGTGATGACGGTGAAAACCTCTGACACATGCAGCTCCCGGAGACGGTCACAG
CTTGTCTGTAAGCGGATGCCGGGAGCAGACAAGCCCGTCAGGGCGCGTCAGCGGGTGTTGGCG
GGTGTCGGGGCTGGCTTAACTATGCGGCATCAGAGCAGATTGTACTGAGAGTGCACCATATGC
GGTGTGAAATACCGCACAGATGCGTAAGGAGAAAATACCGCATCAGGCGCCATTCGCCATTCA
GGCTGCGCAACTGTTGGGAAGGGCGATCGGTGCGGGCCTCTTCGCTATTACGCCAGCTGGCGA
AAGGGGGATGTGCTGCAAGGCGATTAAGTTGGGTAACGCCAGGGTTTTCCCAGTCACGACGT
TGTAAAACGACGGCCAGTAGATCTACGCCAAGCGCGCAATTAACCCTCACTAAAGGGAACAA
AAGCTGGAGCTCCGGATGCAAGGGTTCGAATCCCTTAGCTCTCATTATTTTTGCTTTTTCTC
TTGAGGTCACATGATCGCAAAATGGCAAATGGCACGTGAAGCTGTCGATATTGGGGAACTGT
GGTGGTTGGCAAATGACTAATTAAGTTAGTCAAGGCGCCATCCTCATGAAAACTGTGTAACA
TAATAACCGAAGTGTCGAAAAGGTGGCACCTTGTCCAATTGAACACGCTCGATGAAAAAAAT
AAGATATATATAAGGTTAAGTAAAGCGTCTGTTAGAAAGGAAGTTTTTCCTTTTTCTTGCTC
TCTTGTCTTTTCATCTACTATTTCCTTCGTGTAATACAGGGTCGTCAGATACATAGATACAAT
TCTATTACCCCCATCCATACTCTAGAACTAGTGGATCCCCCGGGCTGCAGGAATTCGATATCA
AGCTTATCGATACCGTCGACCTCGACATGAATGGCGCTATTGGCGGTGACCTATTATTAAATT
TTCCAGATATGTCTGTCCTAGAAAGACAACGTGCCCATTTAAAATACTTAAATCCTACATTCG
ATTCGCCACTTGCGGGTTTCTTTGCTGATAGCAGCATGATAACAGGCGGTGAAATGGATAGT
TATTTATCCACAGCAGGTCTAAACCTACCCATGATGTATGGGGAAACGACAGTTGAGGGCGA
TTCTCGTTTGTCAATCTCCCCAGAGACGACCCTTGGCACAGGTAATTTCAAAAAAAGAAAATT
TGATACTGAAACCAAGGACTGCAATGAGAAGAAAAAAAAAATGACAATGAATAGGGATGATT
TAGTAGAAGAGGGAGAGGAGGAAAAATCCAAAATTACTGAACAAAACAATGGTTCTACCAAG
TCGATTAAGAAAATGAAGCATAAGGCAAAAAAAGAAGAGAACAATTTTTCAAATGATTCATC
AAAGGTTACTAAAGAATTAGAAAAGACAGATTACATTCACGTTAGAGCACGTAGGGGACAGG
CTACTGACTCTCATTCCATTGCGGAGAGAGTCAGGCGTGAAAAAATCTCAGAAAGAATGAAG
TTCTTACAGGACCTAGTTCCTGGGTGTGACAAGATTACGGGTAAAGCTGGTATGTTAGACGA
GATCATCAACTATGTTCAAAGTCTTCAAAGGCAGATTGAATTCTTATCTATGAAGCTGGCTA
TCGTAAATCCTAGGCCAGATTTCGATATGGATGATATTTTCGCAAAAGAAGTCGCAAGTACC
CCAATGACTGTGGTACCCAGCCCAGAAATGGTGTTGTCTGGTTATTCACACGAAATGGTTCAC
AGTGGGTACAGCTCAGAGATGGTGAATTCTGGCTACCTTCATGTTAACCCAATGCAACAAGT
GAATACGTCCTCTGATCCATTATCCTGTTTTAATAATGGTGAAGCGCCAAGTATGTGGGATTC
TCATGTTCAAAATCTATATGGAAATTTGGGAGTAGGTGGTGGGGGTAGCGGAGGCGGTGGTA
GCGGTGGTGGTGGAAGGCGACCAAGTGACAGCAATGCTGTTTCACTGGTTATGCGGCGGATCC
GAAAAGAAAACGTTGATGCCGGTGAACGTGCAAAACAGGCTCTAGCGTTCAGATCTGGCGTA
ATCATGGTCATAGCTGTTTCCTGTGTGAAATTGTTATCCGCTCACAATTCCACACAACATACG
AGCCGGAAGCATAAAGTGTAAAGCCTGGGGTGCCTAATGAGTGAGCTAACTCACATTAATTG
CGTTGCGCTCACTGCCCGCTTTCCAGTCGGGAAACCTGTCGTGCCAGCTGCATTAATGAATCG
GCCAACGCGCGGGGAGAGGCGGTTTGCGTATTGGGCGCTCTTCCGCTTCCTCGCTCACTGACT
CGCTGCGCTCGGTCGTTCGGCTGCGGCGAGCGGTATCAGCTCACTCAAAGGCGGTAATACGGT
TATCCACAGAATCAGGGGATAACGCAGGAAAGAACATGTGAGCAAAAGGCCAGCAAAAGGCC
AGGAACCGTAAAAAGGCCGCGTTGCTGGCGTTTTTCCATAGGCTCCGCCCCCCTGACGAGCAT
CACAAAAATCGACGCTCAAGTCAGAGGTGGCGAAACCCGACAGGACTATAAAGATACCAGGC
GTTTCCCCCTGGAAGCTCCCTCGTGCGCTCTCCTGTTCCGACCCTGCCGCTTACCGGATACCTG
TCCGCCTTTCTCCCTTCGGGAAGCGTGGCGCTTTCTCATAGCTCACGCTGTAGGTATCTCAGT
TCGGTGTAGGTCGTTCGCTCCAAGCTGGGCTGTGTGCACGAACCCCCCGTTCAGCCCGACCGC
TGCGCCTTATCCGGTAACTATCGTCTTGAGTCCAACCCGGTAAGACACGACTTATCGCCACTG
GCAGCAGCCACTGGTAACAGGATTAGCAGAGCGAGGTATGTAGGCGGTGCTACAGAGTTCTT
GAAGTGGTGGCCTAACTACGGCTACACTAGAAGAACAGTATTTGGTATCTGCGCTCTGCTGAA

GCCAGTTACCTTCGGAAAAAGAGTTGGTAGCTCTTGATCCGGCAAACAAACCACCGCTGGTAG
CGGTGGTTTTTTTGTTTGCAAGCAGCAGATTACGCGCAGAAAAAAAGGATCTCAAGAAGATC
CTTTGATCTTTTCTACGGGGTCTGACGCTCAGTGGAACGAAAACTCACGTTAAGGGATTTTGG
TCATGAGATTATCAAAAAGGATCTTCACCTAGATCCTTTTAAATTAAAAATGAAGTTTTAAA
TCAATCTAAAGTATATATGAGTAAACTTGGTCTGACAGTTACCAATGCTTAATCAGTGAGGC
ACCTATCTCAGCGATCTGTCTATTTCGTTCATCCATAGTTGCCTGACTCCCCGTCGTGTAGAT
AACTACGATACGGGAGGGCTTACCATCTGGCCCCAGTGCTGCAATGATACCGCGAGACCCACG
CTCACCGGCTCCAGATTTATCAGCAATAAACCAGCCAGCCGGAAGGGCCGAGCGCAGAAGTGG
TCCTGCAACTTTATCCGCCTCCATCCAGTCTATTAATTGTTGCCGGGAAGCTAGAGTAAGTAG
TTCGCCAGTTAATAGTTTGCGCAACGTTGTTGCCATTGCTACAGGCATCGTGGTGTCACGCTC
GTCGTTTGGTATGGCTTCATTCAGCTCCGGTTCCCAACGATCAAGGCGAGTTACATGATCCCC
CATGTTGTGCAAAAAAGCGGTTAGCTCCTTCGGTCCTCCGATCGTTGTCAGAAGTAAGTTGGC
CGCAGTGTTATCACTCATGGTTATGGCAGCACTGCATAATTCTCTTACTGTCATGCCATCCGT
AAGATGCTTTTCTGTGACTGGTGAGTACTCAACCAAGTCATTCTGAGAATAGTGTATGCGGC
GACCGAGTTGCTCTTGCCCGGCGTCAATACGGGATAATACCGCGCCACATAGCAGAACTTTAA
AAGTGCTCATCATTGGAAAACGTTCTTCGGGGCGAAAACTCTCAAGGATCTTACCGCTGTTGA
GATCCAGTTCGATGTAACCCACTCGTGCACCCAACTGATCTTCAGCATCTTTTACTTTCACCA
GCGTTTCTGGGTGAGCAAAAACAGGAAGGCAAAATGCCGCAAAAAAGGGAATAAGGGCGACA
CGGAAATGTTGAATACTCATACTCTTCCTTTTTCAATATTATTGAAGCATTTATCAGGGTTAT
TGTCTCATGAGCGGATACATATTTGAATGTATTTAGAAAAATAAACAAATAGGGGTTCCGCG
CACATTTCCCCGAAAAGTGCCACCTGACGTCTAAGAAACCATTATTATCATGACATTAACCTA
TAAAAATAGGCGTATCACGAGGCCCTTTCGTC

>CRY2CreNter
TCGCGCGTTTCGGTGATGACGGTGAAAACCTCTGACACATGCAGCTCCCGGAGACGGTCACAG
CTTGTCTGTAAGCGGATGCCGGGAGCAGACAAGCCCGTCAGGGCGCGTCAGCGGGTGTTGGCG
GGTGTCGGGGCTGGCTTAACTATGCGGCATCAGAGCAGATTGTACTGAGAGTGCACCATATGC
GGTGTGAAATACCGCACAGATGCGTAAGGAGAAAATACCGCATCAGGCGCCATTCGCCATTCA
GGCTGCGCAACTGTTGGGAAGGGCGATCGGTGCGGGCCTCTTCGCTATTACGCCAGCTGGCGA
AAGGGGGATGTGCTGCAAGGCGATTAAGTTGGGTAACGCCAGGGTTTTCCCAGTCACGACGT
TGTAAAACGACGGCCAGTAGATCTCAGATACATAGATACAATTCTATTACCCCCATCCATACT
CTAGAACTAGTGGATCCCCCGGGCTGCAGGAATTCGATATCAAGCTTATCGATACCGTCGACC
TCGACATGAAGATGGATAAGAAGACAATCGTTTGGTTCAGAAGGGACTTAAGAATCGAAGAC
AATCCCGCGTTGGCGGCCGCCGCACATGAGGGATCCGTATTTCCGGTGTTTATCTGGTGCCCA
GAAGAAGAAGGGCAATTCTACCCTGGTCGTGCTAGTAGGTGGTGGATGAAGCAATCATTAGC
CCACCTCTCCCAATCACTTAAAGCACTGGGCTCTGATCTCACATTGATTAAGACACATAATAC
GATCTCAGCGATCCTCGACTGCATCAGAGTAACCGGCGCTACAAAGGTTGTGTTCAATCACTT
ATACGACCCTGTCTCATTGGTCAGAGATCATACCGTAAAAGAGAAACTCGTGGAAAGAGGAA
TTAGCGTTCAGTCTTATAACGGCGATCTATTGTATGAACCATGGGAAATATACTGTGAAAAA
GGGAAGCCATTTACTTCATTTAACTCATATTGGAAAAAATGCCTAGATATGTCTATCGAATC
GGTCATGTTGCCTCCCCCCTGGAGGCTAATGCCAATAACCGCAGCTGCTGAAGCTATTTGGGC
GTGCAGTATTGAAGAATTGGGGTTAGAAAATGAAGCTGAAAAACCTTCCAATGCATTACTCA
CAAGAGCTTGGAGTCCCGGATGGTCAAATGCAGATAAATTACTGAATGAATTTATCGAGAAA
CAATTGATTGATTACGCGAAGAATTCCAAAAAAGTCGTGGGAAACTCTACCAGTTTGCTTTCC
CCATATCTCCATTTCGGCGAGATTTCAGTCAGACATGTATTTCAGTGTGCAAGAATGAAACAG
ATCATTTGGGCAAGGGATAAAAACTCGGAAGGTGAAGAATCTGCTGATCTTTTCTTAAGGGG
TATTGGTTTGAGAGAATACTCGCGTTACATCTGCTTCAACTTTCCATTTACGCACGAACAATC
ATTGTTATCGCATCTGAGGTTTTTCCCATGGGATGCAGATGTCGATAAGTTTAAAGCTTGGA
GACAAGGGAGGACAGGTTATCCTCTGGTTGATGCAGGCATGCGTGAATTTTGGGCTACAGGT

TGGATGCACAACAGGATAAGGGTTATCGTGTCATCATTCGCCGTAAAGTTTCTTTTACTGCCG
TGGAAGTGGGGGATGAAGTACTTTTGGGATACATTGCTTGATGCTGACCTCGAGTGCGACAT
ACTAGGATGGCAGTATATTTCTGGTAGTATTCCAGATGGTCATGAACTTGATCGTTTAGATA
ATCCCGCTTTGCAGGGTGCCAAATATGATCCAGAAGGTGAATACATTAGGCAGTGGTTACCTG
AGTTGGCTCGGCTGCCTACTGAATGGATACACCATCCCTGGGATGCTCCTTTAACTGTCCTGA
AAGCATCAGGTGTCGAATTAGGTACCAATTACGCGAAACCTATTGTGGACATCGACACTGCA
AGAGAACTCTTGGCCAAAGCGATATCCAGGACACGAGAAGCGCAGATCATGATCGGGGCCGC
TCCAGATGAAATCGTTGCCGACTCTTTTGAAGCATTAGGCGCCAATACTATTAAAGAACCAGG
GCTATGTCCCAGCGTATCCTCCAATGACCAACAGGTACCATCTGCTGTAAGATATAATGGTTC
AAAAAGAGTTAAACCCGAAGAAGAAGAAGAGCGCGACATGAAGAAAAGCAGAGGCTTTGACG
AAAGAGAACTATTCTCTACCGCGGAAAGTTCCAGTTCCTCCTCAGTCTTTTTCGTTTCGCAGT
CCTGTTCTTTAGCATCTGAAGGTAAGAACCTTGAGGGAATTCAAGATTCTTCTGATCAAATCA
CCACTTCCCTGGGCAAAAATGGTTGCAAGGGCGGTGGAGGGTCCGGTGGGGGTGGAAGCGGA
GGAGGTGGTAGGACGAGTGATGAGGTTCGCAAGAACCTGATGGACATGTTCAGGGATCGCCA
GGCGTTTTCTGAGCATACCTGGAAAATGCTTCTGTCCGTTTGCCGGTCGTGGGCGGCATGGTG
CAAGTTGAATAACCGGAAATGGTTTCCCGCAGAACCTGAAGATGTTCGCGATTATCTTCTATA
TCTTCAGGCGCGCGGTCTGGCAGTAAAAACTATCCAGCAACATTTGGGCCAGCTAAACATGCT
TCATCGTCGGTCCGGGCTGTAGCCATTAACGCGTAAATGATTGCTATAATTATTTGATATTTA
TGGTGACATATGAGAAAGGATTTCAACATCGACGGAAAATATGTAGTGCTGTCTGTAAAGAT
CTGGCGTAATCATGGTCATAGCTGTTTCCTGTGTGAAATTGTTATCCGCTCACAATTCCACAC
AACATACGAGCCGGAAGCATAAAGTGTAAAGCCTGGGGTGCCTAATGAGTGAGCTAACTCAC
ATTAATTGCGTTGCGCTCACTGCCCGCTTTCCAGTCGGGAAACCTGTCGTGCCAGCTGCATTA
ATGAATCGGCCAACGCGCGGGGAGAGGCGGTTTGCGTATTGGGCGCTCTTCCGCTTCCTCGCT
CACTGACTCGCTGCGCTCGGTCGTTCGGCTGCGGCGAGCGGTATCAGCTCACTCAAAGGCGGT
AATACGGTTATCCACAGAATCAGGGGATAACGCAGGAAAGAACATGTGAGCAAAAGGCCAGC
AAAAGGCCAGGAACCGTAAAAAGGCCGCGTTGCTGGCGTTTTTCCATAGGCTCCGCCCCCCTG
ACGAGCATCACAAAAATCGACGCTCAAGTCAGAGGTGGCGAAACCCGACAGGACTATAAAGA
TACCAGGCGTTTCCCCCTGGAAGCTCCCTCGTGCGCTCTCCTGTTCCGACCCTGCCGCTTACCG
GATACCTGTCCGCCTTTCTCCCTTCGGGAAGCGTGGCGCTTTCTCATAGCTCACGCTGTAGGT
ATCTCAGTTCGGTGTAGGTCGTTCGCTCCAAGCTGGGCTGTGTGCACGAACCCCCCGTTCAGC
CCGACCGCTGCGCCTTATCCGGTAACTATCGTCTTGAGTCCAACCCGGTAAGACACGACTTAT
CGCCACTGGCAGCAGCCACTGGTAACAGGATTAGCAGAGCGAGGTATGTAGGCGGTGCTACAG
AGTTCTTGAAGTGGTGGCCTAACTACGGCTACACTAGAAGAACAGTATTTGGTATCTGCGCTC
TGCTGAAGCCAGTTACCTTCGGAAAAAGAGTTGGTAGCTCTTGATCCGGCAAACAAACCACCG
CTGGTAGCGGTGGTTTTTTTGTTTGCAAGCAGCAGATTACGCGCAGAAAAAAAGGATCTCAA
GAAGATCCTTTGATCTTTTCTACGGGGTCTGACGCTCAGTGGAACGAAAACTCACGTTAAGGG
ATTTTGGTCATGAGATTATCAAAAAGGATCTTCACCTAGATCCTTTTAAATTAAAAATGAAG
TTTTAAATCAATCTAAAGTATATATGAGTAAACTTGGTCTGACAGTTACCAATGCTTAATCA
GTGAGGCACCTATCTCAGCGATCTGTCTATTTCGTTCATCCATAGTTGCCTGACTCCCCGTCG
TGTAGATAACTACGATACGGGAGGGCTTACCATCTGGCCCCAGTGCTGCAATGATACCGCGAG
ACCCACGCTCACCGGCTCCAGATTTATCAGCAATAAACCAGCCAGCCGGAAGGGCCGAGCGCA
GAAGTGGTCCTGCAACTTTATCCGCCTCCATCCAGTCTATTAATTGTTGCCGGGAAGCTAGAG
TAAGTAGTTCGCCAGTTAATAGTTTGCGCAACGTTGTTGCCATTGCTACAGGCATCGTGGTGT
CACGCTCGTCGTTTGGTATGGCTTCATTCAGCTCCGGTTCCCAACGATCAAGGCGAGTTACAT
GATCCCCCATGTTGTGCAAAAAAGCGGTTAGCTCCTTCGGTCCTCCGATCGTTGTCAGAAGTA
AGTTGGCCGCAGTGTTATCACTCATGGTTATGGCAGCACTGCATAATTCTCTTACTGTCATGC
CATCCGTAAGATGCTTTTCTGTGACTGGTGAGTACTCAACCAAGTCATTCTGAGAATAGTGTA
TGCGGCGACCGAGTTGCTCTTGCCCGGCGTCAATACGGGATAATACCGCGCCACATAGCAGAA
CTTTAAAAGTGCTCATCATTGGAAAACGTTCTTCGGGGCGAAAACTCTCAAGGATCTTACCGC

TGTTGAGATCCAGTTCGATGTAACCCACTCGTGCACCCAACTGATCTTCAGCATCTTTTACTT
TCACCAGCGTTTCTGGGTGAGCAAAAACAGGAAGGCAAAATGCCGCAAAAAAGGGAATAAGG
GCGACACGGAAATGTTGAATACTCATACTCTTCCTTTTTCAATATTATTGAAGCATTTATCAG
GGTTATTGTCTCATGAGCGGATACATATTTGAATGTATTTAGAAAAATAAACAAATAGGGGT
TCCGCGCACATTTCCCCGAAAAGTGCCACCTGACGTCTAAGAAACCATTATTATCATGACATT
AACCTATAAAAATAGGCGTATCACGAGGCCCTTTCGTC

>gERG9
ggatcctagactctcgaggcgaatttcttatgatttatgatttttattattaaataagttataaaaaaaataagtgtatacaaatttt
aaagtgactcttaggttttaaaacgaaaattcttattcttgagtaactctttcctgtaggtcaggttgctttctcaggtatagcatga
ggtcgctcttattgaccacacctctaccggcatgccgagcaaatgcctgcaaatcgctccccatttctctagagcggccgtggtat
cgtttagattggcaattacagACgtcttagctcacatgcttataactaattacatgactcgaagacataaaaaacaaaaaaagc
accaccgactcggtgccactttttcaagttgataacggactagccttattttaacttgctatttCTAGCTCTAAAACATTG
TAATAGCTTTCCCATtgatcatttatctttcactgcggagaagtttcgaacgccgaaacatgcgcaccaactttcacttct
acagcgtttgaccaaaatcttttgaacagaacattgtagggtgtgaaaaaatgcgcacctttaccgctagc

Supplementary Text S2: Peptide sequences.

>LiCre (from pGY466)
MRGSHHHHHHGSLATTLERIEKNFVITDPRLPDNPIIFASDSFLQLTEYSREEILGRNCRFLQGP
ETDRATVRKIRDAIDNQTEVTVQLINYTKSGKKFWNVFHLQPMRDYKGDVQYFIGVQLDGTE
RLHGAAEREAVCLIKKTAFQIDSDEVRKNLMDMFRDRQAFSEHTWKMLLSVCRSWAAWCKL
NNRKWFPAEPEDVRDYLLYLQARGLAVKTIQQHLGQLNMLHRRSGLPRPSDSNAVSLVMRRI
RKENVDAGERAKQALAFERTDFDQVRSLMENSDRCQDIRNLAFLGIAYNTLLRIAEIARIRVKD
ISRTDGGRMLIHIGRTKTLVSTAGVEKALSLGVTKLVERWISVSGVADDPNNYLFCRVRKNGVA
APSATSQLSTRALEGIFEATHRLIYGAKDDSGQRYLAWSGHSARVGAARDMARAGVSIPEIMQ
AGGWTNVNIVMNYIRNLDSETGAMVRLLAAGD

>LOV2_Cre32 (from pGY415)
MRGSHHHHHHGSLATTLERIEKNFVITDPRLPDNPIIFASDSFLQLTEYSREEILGRNCRFLQGP
ETDRATVRKIRDAIDNQTEVTVQLINYTKSGKKFWNVFHLQPMRDYKGDVQYFIGVQLDGTE
RLHGAAEREAVCLIKKTAFQIARDRQAFSEHTWKMLLSVCRSWAAWCKLNNRKWFPAEPED
VRDYLLYLQARGLAVKTIQQHLGQLNMLHRRSGLPRPSDSNAVSLVMRRIRKENVDAGERAK
QALAFERTDFDQVRSLMENSDRCQDIRNLAFLGIAYNTLLRIAEIARIRVKDISRTDGGRMLIHI
GRTKTLVSTAGVEKALSLGVTKLVERWISVSGVADDPNNYLFCRVRKNGVAAPSATSQLSTRA
LEGIFEATHRLIYGAKDDSGQRYLAWSGHSARVGAARDMARAGVSIPEIMQAGGWTNVNIVM
NYIRNLDSETGAMVRLLEDGD

> NLS-pMag-CreC60-343 (from pGY491)
MVPKKKRKVGGHTLYAPGGYDIMGYLRQIRNRPNPQVELGPVDTSCALILCDLKQKDTPIVYAS
EAFLYMTGYSNAEVLGRNCRFLQSPDGMVKPKSTRKYVDSNTINTMRKAIDRNAEVQVEVVN
FKKNGQRFVNFLTMIPVRDETGEYRYSMGFQCETEGTNRKWFPAEPEDVRDYLLYLQARGLA
VKTIQQHLGQLNMLHRRSGLPRPSDSNAVSLVMRRIRKENVDAGERAKQALAFERTDFDQVR
SLMENSDRCQDIRNLAFLGIAYNTLLRIAEIARIRVKDISRTDGGRMLIHIGRTKTLVSTAGVEK
ALSLGVTKLVERWISVSGVADDPNNYLFCRVRKNGVAAPSATSQLSTRALEGIFEATHRLIYGA
KDDSGQRYLAWSGHSARVGAARDMARAGVSIPEIMQAGGWTNVNIVMNYIRNLDSETGAMV
RLLEDGD

> CreN18-59-nMag-NLS (from pGY501)
MATSDEVRKNLMDMFRDRQAFSEHTWKMLLSVCRSWAAWCKLNGTHTLYAPGGYDIMGYL
DQIGNRPNPQVELGPVDTSCALILCDLKQKDTPIVYASEAFLYMTGYSNAEVLGRNCRFLQSPD
GMVKPKSTRKYVDSNTINTMRKAIDRNAEVQVEVVNFKKNGQRFVNFLTMIPVRDETGEYRY
SMGFQCETEGGSGGVPKKKRKV

> CRY2L348F-CreN19-104 (from pGY531)
MKMDKKTIVWFRRDLRIEDNPALAAAAHEGSVFPVFIWCPEEEGQFYPGRASRWWMKQSLA
HLSQSLKALGSDLTLIKTHNTISAILDCIRVTGATKVVFNHLYDPVSLVRDHTVKEKLVERGISV
QSYNGDLLYEPWEIYCEKGKPFTSFNSYWKKCLDMSIESVMLPPPWRLMPITAAAEAIWACSI
EELGLENEAEKPSNALLTRAWSPGWSNADKLLNEFIEKQLIDYAKNSKKVVGNSTSLLSPYLHF
GEISVRHVFQCARMKQIIWARDKNSEGEESADLFLRGIGLREYSRYICFNFPFTHEQSLLSHLRF
FPWDADVDKFKAWRQGRTGYPLVDAGMREFWATGWMHNRIRVIVSSFAVKFLLLPWKWG
MKYFWDTLLDADLECDILGWQYISGSIPDGHELDRLDNPALQGAKYDPEGEYIRQWLPELARL
PTEWIHHPWDAPLTVLKASGVELGTNYAKPIVDIDTARELLAKAISRTREAQIMIGAAPDEIVA

DSFEALGANTIKEPGLCPSVSSNDQQVPSAVRYNGSKRVKPEEEEERDMKKSRGFDERELFSTA
ESSSSSSVFFVSQSCSLASEGKNLEGIQDSSDQITTSLGKNGCKGGGGSGGGGSGGGGRTSDEVRK
NLMDMFRDRQAFSEHTWKMLLSVCRSWAAWCKLNNRKWFPAEPEDVRDYLLYLQARGLAV
KTIQQHLGQLNMLHRRSGL

> CIB1-CreC106-343 (from pGY532)
MNGAIGGDLLLNFPDMSVLERQRAHLKYLNPTFDSPLAGFFADSSMITGGEMDSYLSTAGLNL
PMMYGETTVEGDSRLSISPETTLGTGNFKKRKFDTETKDCNEKKKKMTMNRDDLVEEGEEEK
SKITEQNNGSTKSIKKMKHKAKKEENNFSNDSSKVTKELEKTDYIHVRARRGQATDSHSIAER
VRREKISERMKFLQDLVPGCDKITGKAGMLDEIINYVQSLQRQIEFLSMKLAIVNPRPDFDMDD
IFAKEVASTPMTVVPSPEMVLSGYSHEMVHSGYSSEMVNSGYLHVNPMQQVNTSSDPLSCFNN
GEAPSMWDSHVQNLYGNLGVGGGGSGGGGSGGGGRRPSDSNAVSLVMRRIRKENVDAGERAK
QALAFERTDFDQVRSLMENSDRCQDIRNLAFLGIAYNTLLRIAEIARIRVKDISRTDGGRMLIHI
GRTKTLVSTAGVEKALSLGVTKLVERWISVSGVADDPNNYLFCRVRKNGVAAPSATSQLSTRA
LEGIFEATHRLIYGAKDDSGQRYLAWSGHSARVGAARDMARAGVSIPEIMQAGGWTNVNIVM
NYIRNLDSETGAMVRLLEDGD
